# Tracking biological hallucinations in single-cell perturbation predictions using scArchon, a comprehensive benchmarking platform

**DOI:** 10.1101/2025.06.23.661046

**Authors:** Jean Radig, Robin Droit, Daria Doncevic, Albert Li, Duc Thien Bui, Luis Herfurth, Thaddeus Kühn, Carl Herrmann

**Affiliations:** Institute of Pharmacy and Molecular Biotechnology (IPMB), Heidelberg University; BioQuant, Heidelberg University

## Abstract

The accurate prediction of cellular responses to perturbations, such as drug treatments, remains a pivotal challenge in single-cell transcriptomics. While numerous deep learning tools have been developed for this task, their systematic benchmarking across diverse datasets and performance metrics has been limited. Here, we present scArchon, a reproducible, modular benchmarking platform built on Snakemake, designed to evaluate perturbation response prediction tools in an unbiased and extensible manner. Employing four representative single-cell RNA-seq datasets, we compare leading methods such as scGen, CPA, trVAE, scPRAM, scVIDR, scDisInFact, SCREEN, scPreGAN, and CellOT against baselines. We assess model performance using a composite of statistical and biological metrics. Our analysis reveals heterogeneous performance: while methods like trVAE, scGen, scPRAM, and scVIDR achieve robust results across multiple datasets, other tools occasionally underperform even compared to linear or control baselines. Notably, models with favorable quantitative scores may fail to retain key biological perturbation signatures, underscoring the need for gene-level evaluation. scArchon provides a unified, extensible foundation for large-scale, standardized benchmarking of perturbation prediction tools, facilitating methodological transparency and accelerating development in this rapidly evolving field. We encourage adoption of scArchon and sharing of containerized tools to drive progress in single-cell perturbation modeling.

## Introduction

Understanding inter- and intra-patient variability is crucial in personalized medicine, as individual differences in gene expression and cellular composition can significantly influence drug responses (Koumpouros & Yiannis, 2025). Single-cell technologies, such as single-cell RNA sequencing (scRNA-seq), have emerged as powerful tools to dissect this heterogeneity at the individual cell level. By analyzing gene expression variability among different cell types and across individuals, researchers can identify specific cellular subpopulations that may respond differently to molecular perturbations such as therapeutic agents. Integrating single-cell analyses with high-throughput drug screening platforms enables the assessment of drug efficacy and toxicity in diverse cellular contexts, facilitating the development of tailored treatment strategies (Monfort-Lanzas et al., 2025). For instance, studies have demonstrated that combining scRNA-seq with drug sensitivity profiling (Dini et al., 2025) can reveal novel prognostic markers and potential therapeutic targets in cancers like liver cancer, thereby enhancing the precision of treatment approaches (Zhou et al., 2024).

Recent progress in machine learning has introduced new opportunities for in-silico predictions of the transcriptional response of cells subjected to drug treatments or other perturbations (Eckhart et al., 2024). Numerous tools have been developed to predict cellular changes under different stimuli, many claiming to generalize to new, unseen cells (out-of-distribution problem) (Sharifi-Noghabi et al., 2021). However, newer methods often neglect benchmarking against all earlier tools, resulting in fragmented and biased comparisons (Fig 1A). Moreover, inconsistent use of performance metrics complicates overarching evaluations. So far, no external comprehensive and reproducible benchmark study of perturbation prediction tools has been conducted. Existing benchmarking efforts neither encompass a comprehensive list of tools nor provide unified, reproducible frameworks (L. Li et al., 2024; Wu et al., 2024; Ziemann et al., 2023).

**Fig 1.**
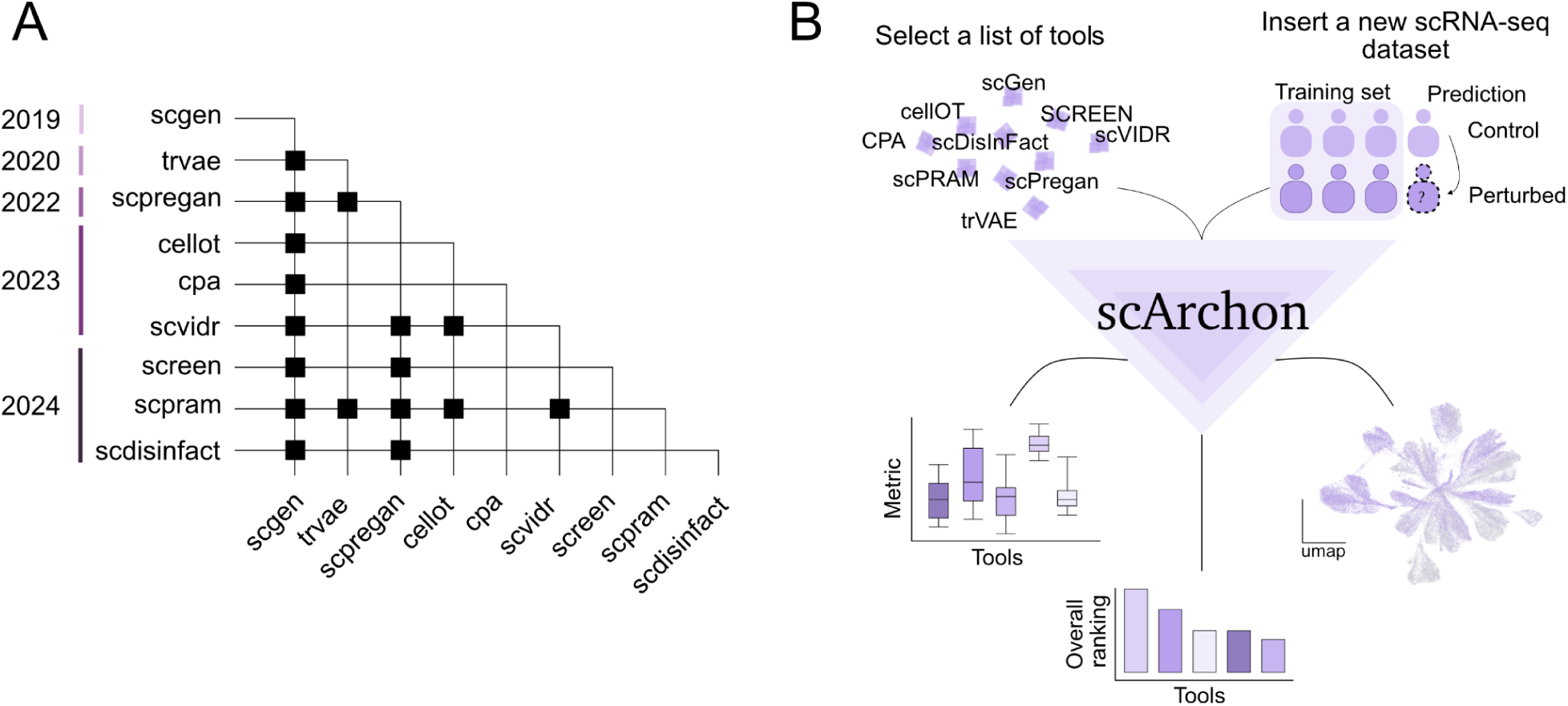
scArchon, a comprehensive benchmarking platform. **A.** Matrix indicating benchmarking comparisons among published tools together with their year of publication. Each row represents a tool that has been benchmarked against the tools listed in the columns; a black dot denotes a reported comparison. **B.** Overview of the scArchon workflow. scArchon generates perturbation predictions of selected tools on user-provided datasets. For each tool, it outputs visual representation of the results as well as quantitative and biological metrics.

To address these issues, we introduce scArchon (Fig 1B), a systematic benchmarking framework for evaluating the latest and most widely used machine learning tools designed to predict drug-induced cellular changes. scArchon standardizes performance evaluation across a curated set of datasets, ensures fair comparisons using consistent metrics, and offers fully reproducible pipelines. Notably, scArchon enables users not only to replicate the results presented here but also to easily apply all tools to their own datasets. Unlike previous benchmarking efforts, scArchon resolves common software environment incompatibility issues by providing containerized, scalable solutions that simplify running and fine-tuning models.

We selected nine tools for benchmarking based on their stated ability to predict drug-induced cellular changes. These tools are: trVAE (Lotfollahi et al., 2020), scDisInFact (Zhang et al., 2024), scVIDR (Kana et al., 2023), scPRAM (Jiang et al., 2024), scGen (Lotfollahi et al., 2019), SCREEN (Wang et al., 2024), CPA (Lotfollahi et al., 2023), scPreGAN (Wei et al., 2022), and CellOT (Bunne et al., 2023). Notably, we limited our study to tools investigating non-genetic perturbations as we consider this to be a separate task compared to predicting the effect of gene knock-outs. Such tools have already been benchmarked previously (Ahlmann-Eltze et al., 2024).

The selected tools differ in their underlying neural network architectures. The most common design involves an encoder that projects the single-cell RNA-sequencing expression matrix into a latent space, where a mapping is learned between the control and perturbed states. In some models, such as scGen, this mapping is linear. In others, including CellOT, scVIDR, scPRAM, and SCREEN, the mapping is based on optimal transport. Alternative architectures are also represented: scPreGAN, for instance, employs a generative adversarial network framework to generate perturbed states from control data. This diversity in modeling approaches allows for a systematic evaluation of how different architectures and assumptions impact performance across various biological scenarios.

The tools were evaluated on four publicly available and widely used datasets: a dataset on interferon treatment of human PBMC cells (called Kang in the remaining article) (Kang et al., 2017), response of mouse epithelial cells to parasite infection (H. Poly) (Haber et al., 2017), response of cells from different species to LPS treatment (Species) (Hagai et al., 2018), and glioblastoma single-cell data from patients treated with panabinostat (Glioblastoma) (Zhao et al., 2021) (Supplementary Figs 1-4), representing increasing prediction challenges. The Kang, H. Poly and Species dataset were chosen specifically because they are curated, preprocessed, and directly accessible online, minimizing variability due to data handling and ensuring fair model comparisons. Besides, they feature clear and well-defined perturbation effects, making them suitable for baseline evaluations. The glioblastoma dataset was selected for its clinical relevance. It presents a more subtle, clinically realistic challenge, enabling us to assess tool robustness under more variable conditions.

By systematically comparing tools on unified tasks and datasets using a variety of metrics, scArchon provides researchers with a clear understanding of each method’s strengths and limitations. Its automated, extensible pipeline ensures that benchmarking remains reproducible, scalable, and relevant as new tools emerge. Beyond identifying the best-performing models, this work emphasizes the importance of testing methods on realistic, challenging datasets to ensure their practical applicability.

## Results

### Experimental design of the benchmark study

We developed scArchon, a benchmarking platform including nine state-of-the-art tools performing perturbation predictions. To ensure that our comparison is fair towards all tools, we first confirmed that we were able to reproduce the tools’ results with our pipeline (Supplementary Notes).

We selected four different datasets of varying complexity. We first define the terms that we will use in this benchmark. For all four datasets, models were trained to predict the *perturbed state* (stimulation, infection or treatment) from the *control state* using all available *covariates* (all cell types, all species or all patients) except one, which was reserved for evaluation. This task is referred to as *out-of-distribution* prediction and tests a model’s ability to generalize to unseen groups. Furthermore, each experiment was repeated in a leave-one-out cross-validation approach iterating over all values of the covariate. We refer to these separate experiments as *prediction folds* throughout the manuscript.

To comprehensively evaluate model performance, we employed complementary approaches: dimensionality reduction visualizations (PCA, t-SNE, UMAP) as well as quantitative metrics (including Mean Squared Error (MSE), Wasserstein distance (Panaretos & Zemel, 2019), and t-tests). We also assessed model performance by comparing differentially expressed genes (DEGs) between control and perturbed cells to those between control and predicted cells, measuring their overlap. Additionally, we performed gene set enrichment analysis on the top DEGs to compare enriched biological terms. For the Kang and H. Poly datasets, we used Gene Ontology (GO) terms (Gene Ontology Consortium et al., 2023), while for the glioblastoma dataset, we used Molecular Signatures Database (MSigDB) cancer hallmark gene sets (Liberzon et al., 2015), which are specifically curated to reflect key oncogenic pathways and are more relevant for cancer-related contexts. This allowed us to evaluate whether the models captured both gene-level changes and broader functional patterns.

### Choice of dimensionality reduction method affects interpretation of results

In the original publications of the different tools, performance is often illustrated through visualization in a reduced-dimensional space, aiming to show the alignment between predicted and perturbed cell states. However, the dimensionality reduction techniques used differ between studies, as certain methods may visually suggest stronger overlap than others. We therefore aimed at systematically showcasing all representations for a fairer comparison between tools.

As a primary example, we present the results obtained on the Kang dataset (Fig 2A–B). This dataset includes seven immune cell types, each with paired control and perturbed conditions. We analyze the prediction fold where models were trained on all cell types except CD4T cells, which were held out as an independent test set (Fig 2A).

**Fig 2.**
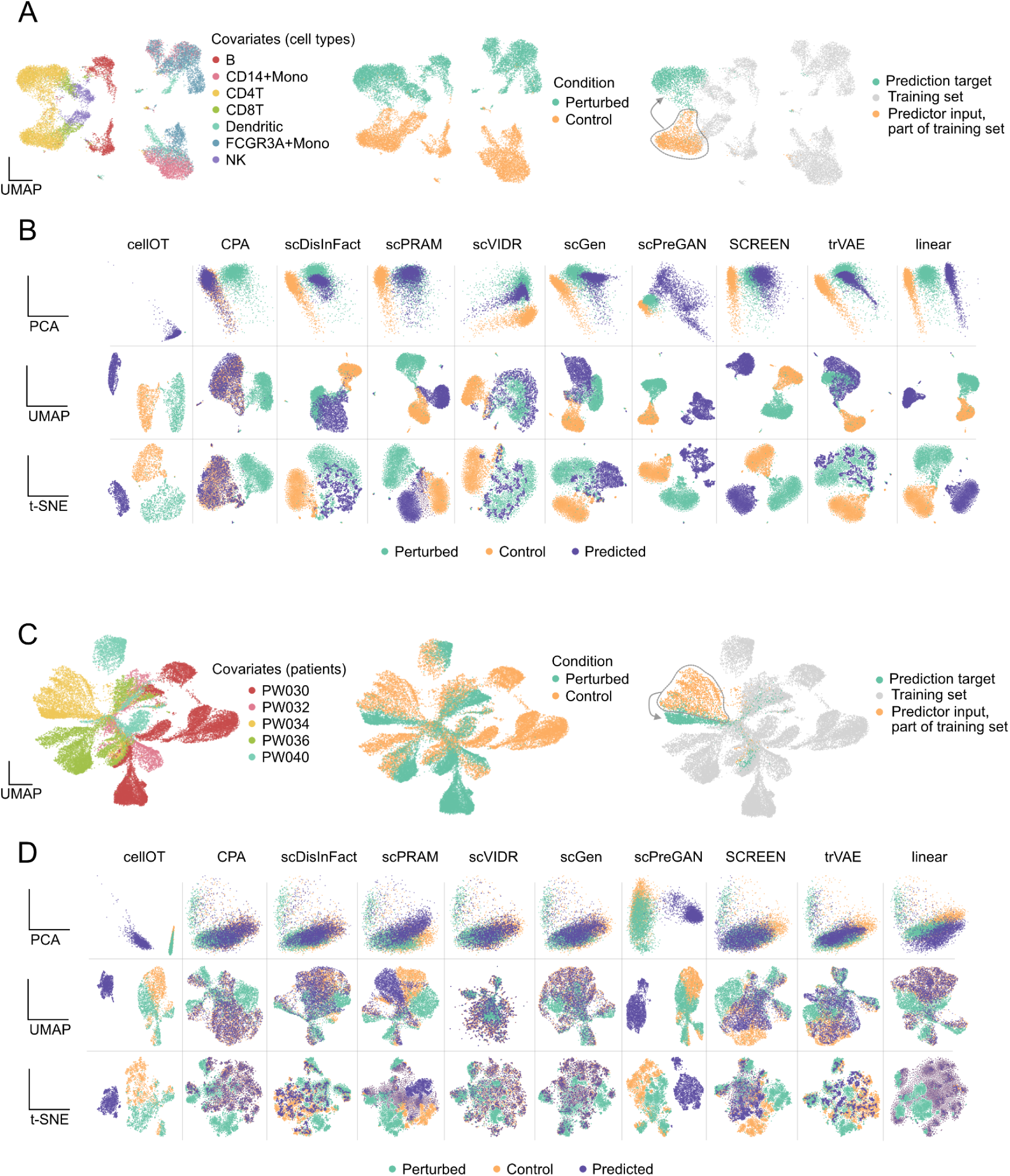
Heterogeneity of the predictions across tools and datasets. The results of CD4T cells from Kang (**A,B**) and the PW034 patient from the Glioblastoma (**C,D**) dataset are shown. **A,C.** UMAP representation of the dataset colored by covariate (left), condition (middle), and experimental setup (right). **B,D.** PCA, UMAP and t-SNE embeddings of control, perturbed and predicted cells for the different tools.

In the PCA representation (Fig 2B), the overlap between predicted and perturbed cells is most evident for scPRAM, scDisInFact, scGen, scVIDR, and trVAE, indicating that these models are able to closely approximate the true perturbation response. In contrast, CellOT fails to achieve any overlap; predicted and perturbed cells are confined to a restricted region of the PCA space. CPA consistently maps predicted cells onto the control distribution rather than the perturbed one. scPreGAN does not produce overlap with either the control or the perturbed cells, indicating a lack of alignment with both reference distributions. The linear model shifts the control cells toward the perturbed state but fails to achieve overlap, which reflects its limited capacity to capture the complexity of the perturbation response.

When examining the UMAP representation, we observe that some methods that appeared to perform well in PCA space show reduced overlap. Notably, scPRAM, which exhibits near-perfect overlap in PCA, shows a clear separation between predicted and perturbed cells in UMAP. A similar trend is observed for scVIDR and SCREEN, where overlap decreases when switching from PCA to UMAP. On the other hand, scDisInFact, scGen, and trVAE produce consistent overlaps across both PCA and UMAP. For the remaining models, UMAP often further emphasizes the lack of overlap already seen in PCA. Finally, the t-SNE representation closely mirrors the patterns seen with UMAP and does not provide additional insights beyond those already captured.

This example highlights the risk of selective visualization. It also underscores the challenge of drawing reliable conclusions from low-dimensional representations. For instance, one might interpret strong alignment between predicted and perturbed cells using PCA, but arrive at the opposite conclusion when viewing the same data through UMAP (Abdi & Williams, 2010; Becht et al., 2018). Such inconsistencies reveal the inherent subjectivity of visual interpretations based on dimensionality reduction.

To illustrate this on a more challenging dataset, we reproduce these results using the glioblastoma dataset (Fig 2C) (Abdi & Williams, 2010; Zhao et al., 2021). Here, the objective is to predict the effect of panobinostat treatment on control cells from patient PW034. In Fig 2D, we observe that even tools that performed well on the Kang dataset tend to produce predicted cells that align more closely with the control group rather than the perturbed group. This pattern suggests a broader limitation in tool generalization when the perturbation effect is masked by confounding effects like inter-patient variability. Results obtained on all experiments for all datasets can be found in Supplementary Figs 5-8.

### Some tools perform no better than a linear model

We quantified tool performance using multiple metrics aggregated over a series of systematic experiments. Results were compared against a *linear model*, which predicts perturbation by adding the mean expression difference between perturbed and control cells to the cells being predicted. The *control baseline* consists simply of the unperturbed cells and represents the absence of any effect.

Tools were ranked according to their average performance across prediction folds for each dataset. The highest mean score for a given metric and dataset received the top rank, with other tools ranked sequentially. These rankings, illustrating each tool’s relative position within individual datasets, are shown in Fig 3A. The top performing tools in the overall ranking appear to recapitulate the visual inspection from the previous section. Looking at individual datasets, we observe heterogeneity in the performance of the tools, for instance scDisInFact performing poorly on H. Poly but well on Species and Glioblastoma.

**Fig 3.**
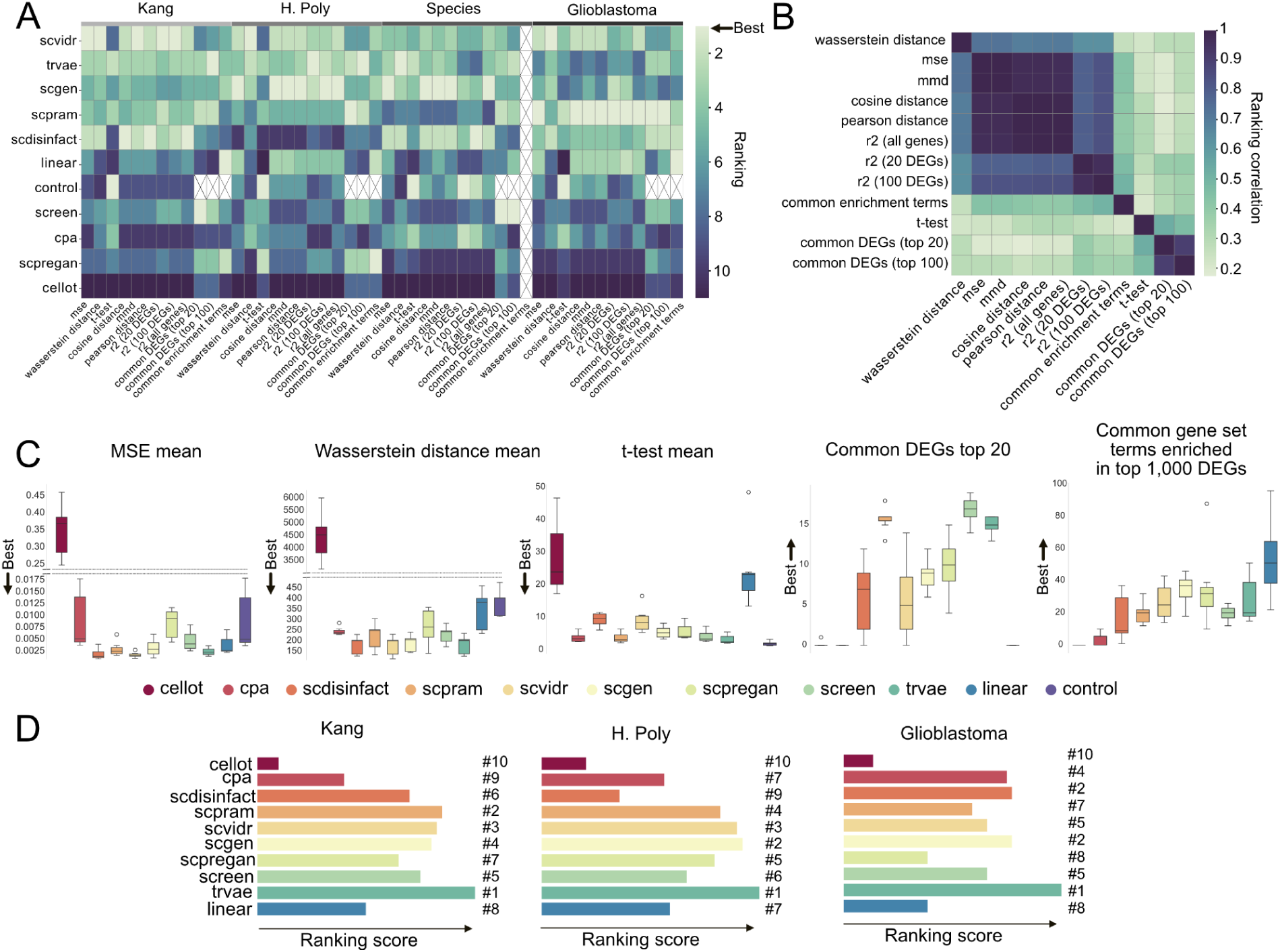
Analysis of scArchon’s output across tools and datasets. **A.** Heatmap showing the performance of the different tools across datasets. Each entry represents the ranking of one tool for a particular metric and dataset. The ranking is based on the average score across all prediction folds (cell types, patients or species) for this dataset. The tools are sorted by their overall ranking. By construction, DEG-based metrics cannot be evaluated on the control baseline (crossed out entries). Common enrichment terms could not be computed for the Species dataset because the genes are not mapped to the corresponding annotation database. **B.** Correlation between tool rankings across different metrics, showing the similarity between different metrics. **C.** Boxplots showing five evaluation metrics computed on the Kang dataset. Each box represents the mean result over the prediction folds. **D.** Overall ranking of each tool across datasets. x-axis represents an ad-hoc ranking score (see Methods). The Species dataset was excluded, as common gene set enrichment could not be computed for this dataset.

Consistent with (Ji et al., 2023), we observed that some metrics produced similar tool rankings (Fig 3A). By definition, some metrics are mathematically related, such as MSE and R2. Indeed, looking at the correlation of the rankings produced by the different metrics, we observe high correlation on some aggregate metrics (Fig 3B). To generate an overall ranking across datasets, avoiding these ranking biases, we selected a subset of metrics that capture distinct performance aspects and are sufficiently uncorrelated, avoiding bias toward tools excelling in only a narrow set of criteria. Using all metrics despite their high correlation could unfairly favor tools optimized for specific measures (L. Li et al., 2024).

After analyzing metric correlations, we selected the following five for our composite evaluation: MSE, Wasserstein distance, t-test score, overlap in top 20 DEGs, and overlap in enriched gene sets derived from the top 1,000 DEGs. This combination balances statistical accuracy with biological interpretability while reducing redundancy. Note that common gene set enrichment could not be computed for the Species dataset because the genes are not mapped to the corresponding annotation database.

To illustrate tool performance, we analysed results on all datasets across the selected metrics (Fig 3C for Kang and Supplementary Figs 5-8 for the others). On the Kang dataset, certain tools consistently ranked high on measures such as MSE, Wasserstein distance, and t-test statistics. Notably, scVIDR, scPRAM, scGen, scDisInFact, and trVAE were among the top performers with low variability across the prediction folds. In contrast, CellOT consistently showed the poorest performance, with CPA and SCREEN also ranking low. Importantly, several tools performed worse than both the linear model and the control baseline, indicating that their predictions deviate further from the true perturbed state than simply using the unaltered control or applying a constant mean shift. This suggests that some models may generate noisy or biologically implausible predictions. More strikingly, considering the t-test metric, which evaluates differences in mean expression relative to variance, no tool outperformed the control baseline.

Interestingly, evaluation based on common DEGs and enriched biological terms revealed that some tools with lower quantitative metric scores still captured meaningful biological signals. For example, SCREEN and scPreGAN ranked among the top performers in these biological assessments. The reverse, however, was also true: tools like CPA or scDisInFact had fair quantitative metric values, however poor overlap at the level of DEGs. These results highlight a key limitation: models that appear convincing in aggregate metrics may fail to preserve the underlying gene-level perturbation structure. This underscores the importance of combining rigorous gene-level quantitative evaluations to accurately interpret and apply perturbation prediction models. Results obtained on the other datasets can be found in Supplementary Figs 6B-8B.

To provide a high-level comparison of tool performance across datasets, we computed a unified ranking based on the five key evaluation metrics shown in Fig 3C. We normalized rankings into a single composite score ranging from 0 to 1, where a score of 1 indicates a tool that consistently ranked first across all experiments and metrics, while a score of 0 indicates a tool that consistently ranked last.

This ranking approach was applied across the Kang, the H. Poly and the Glioblastoma datasets in the benchmark. The Species dataset was excluded because gene set enrichment could not be computed. The results are summarized in Fig 3D. We find that trVAE is the top-performing method overall. Interestingly, scDisInFact, which ranks relatively low on the Kang (6th) and H. poly (9th) datasets, achieves second place on the glioblastoma dataset, a task which is arguably more complex.

The linear model ranks twice in 8th place, and once in 7th. Notably, tools such as CellOT, CPA and scPreGAN tend to sometimes fall below the linear model in performance. In contrast, methods like scGen, scPRAM, and scVIDR consistently rank near the top, demonstrating more robust performance across different biological contexts.

### Deeper analysis of biological metrics reveals hallucinations of the prediction models

A closer inspection of the results in Fig 3C (right panel) reveals that the linear model achieved the highest overlap in enriched GO terms with the perturbed condition on the Kang dataset, with an average of 54 shared terms over the prediction folds. This was followed by scPreGAN and scGen, with 36 and 35 overlapping terms, respectively, while trVAE and scVIDR also performed well, with 29 and 27 shared terms. However, these values must be interpreted with caution, as the number of shared terms is inherently influenced by the total number of predicted enriched terms (referred to as predicted terms). Tools that produce a large number of predicted terms may inflate overlap, increasing the risk of false positives. For instance, although scGen shows strong performance both qualitatively and across multiple metrics (Figs 2B and 3A–D), it predicted 252 enriched GO terms for CD4 T cells, compared to only 72 in the perturbed condition (referred to as baseline terms) (Fig 4A-B). Of these, 41 terms overlapped with the baseline terms.

**Fig 4:**
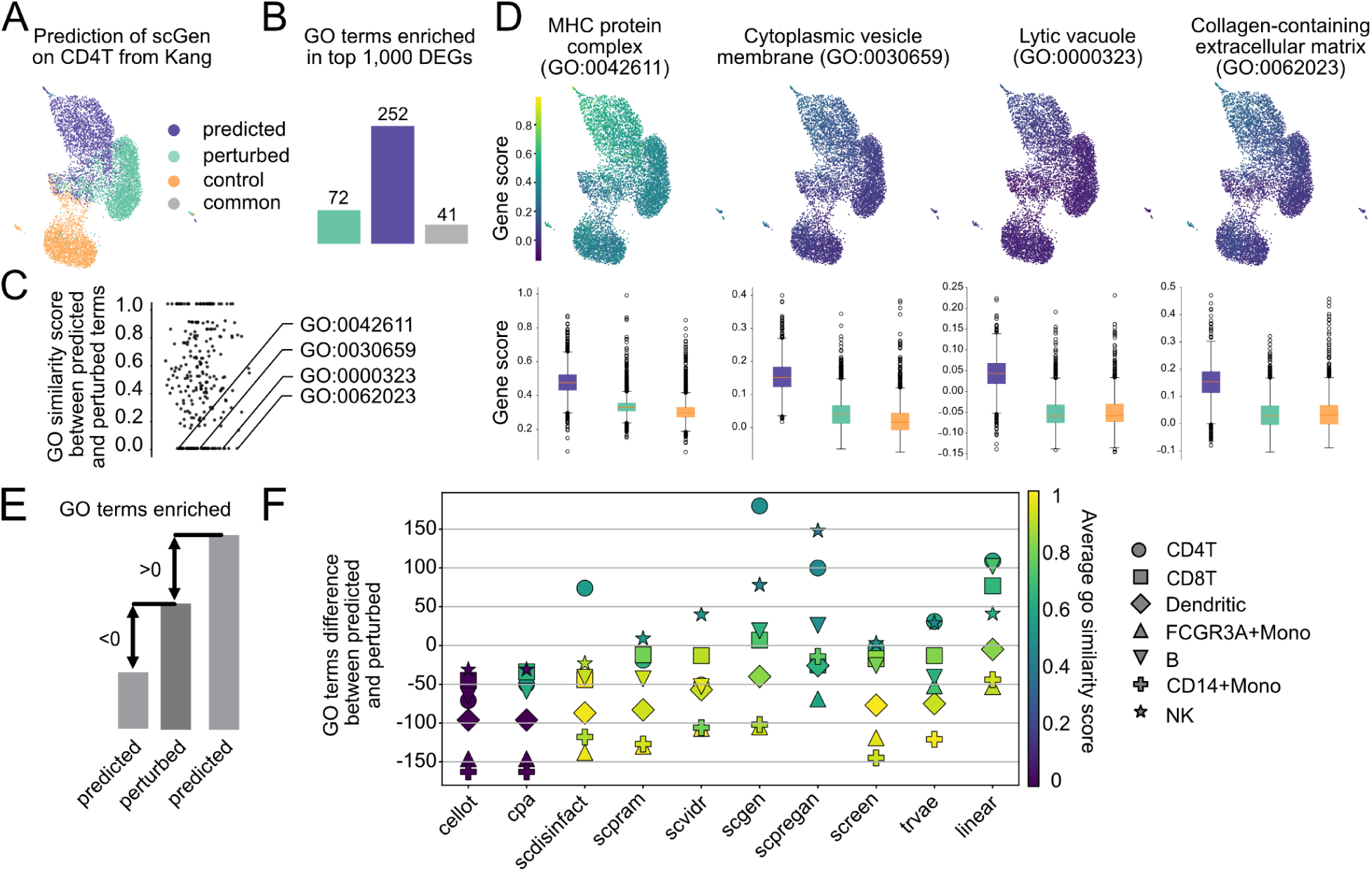
Deeper analysis of biological metrics reveals hallucination of prediction methods. **A.** UMAP showing the predicted expression profile alongside control and perturbed cells. **B.** Number of Gene Ontology (GO) terms enriched among the top 1,000 differentially expressed genes between control and predicted (predicted terms, blue) or control and perturbed cells (baseline terms, green), with grey indicating the shared enriched terms. **C.** Dotplot showing the highest semantic similarity score for each predicted term compared to all baseline terms. Four predicted GO terms with a maximum similarity score of zero were arbitrarily selected for deeper analysis. **D.** Signature scores based on average gene expression for the four selected GO terms with zero similarity. Boxplots show the score differences between predicted and perturbed cells, all of which are statistically significant (Mann–Whitney U test, *p* < 0.01). **E.** Computation schematic. We compare the number of predicted terms with the number of baseline terms. The difference is positive when more GO terms are predicted than in the reference, and negative when fewer are predicted. **F.** Results across experiments. Each dot represents the prediction fold (celltype), the Y axis represents the difference between the number of predicted and baseline terms. The color indicates the average semantic similarity between predicted and baseline terms.

To evaluate whether the additional predicted terms were biologically meaningful, we computed semantic similarity scores between predicted terms and baseline terms (Fig 4C). As expected, the common terms had a similarity score of 1. However, several predicted-only terms had a similarity score of 0 to all baseline terms. To investigate whether these might represent false positives, we arbitrarily selected four such terms and examined the expression of the genes associated with them across control, predicted, and perturbed conditions (Fig 4D). In each case, the expression of the associated genes was significantly higher in the predicted condition than in the perturbed one (based on a Mann-Whitney U test). This suggests that these predictions might result from artificially elevated gene expression, potentially leading to spurious GO term enrichment and misleading downstream biological conclusions.

We performed this analysis on the Kang dataset prediction folds, comparing the number of predicted terms by each method to the number of baseline terms (Fig 4E–F). Some tools, such as scGen and scPreGAN, tend to predict more enriched terms than observed in the perturbed condition, while others consistently yield fewer. Interestingly, tools that predict fewer terms than the ground truth often achieve higher average semantic similarity scores—indicating fewer, but more biologically relevant predictions (for example SCREEN on monocytes). However, scGen and scPreGAN do not always over-predict. As shown in Supplementary Fig 9 for the H. poly dataset (also evaluated using GO similarity), these tools often predict a similar or even smaller number of terms compared to others.

These findings highlight that biological evaluation of prediction results can be challenging, as tools may produce misleading biological enrichments, so-called hallucinations. High overlap in enriched terms or DEGs does not always reflect true biological relevance, especially when driven by inflated or unstable predictions. This underscores the need for careful interpretation and complementary analyses when assessing the biological fidelity of perturbation models.

### Tools have different robustness in ablation experiments

In addition to evaluating model performance, we examined tool efficiency and robustness through runtime and ablation studies. Fig 5A shows the average runtime per prediction fold across datasets of increasing cell numbers. All tools scale with dataset size except CellOT, which maintains a constant—and highest overall—computation time. To test robustness, we conducted an ablation experiment by progressively reducing the number of perturbed cells to 80%, 60%, and 40% (Fig 5B). Most tools remained stable on MSE and Wasserstein scores, with slight worsening of performance observed for SCREEN, scPreGAN, and CellOT (Fig 5C). For DEG overlap, scDisInFact showed minor fluctuations but remained consistent, and similar patterns were seen across other tools. A general decline in the number of enriched terms was noted with fewer perturbed cells.

**Fig 5:**
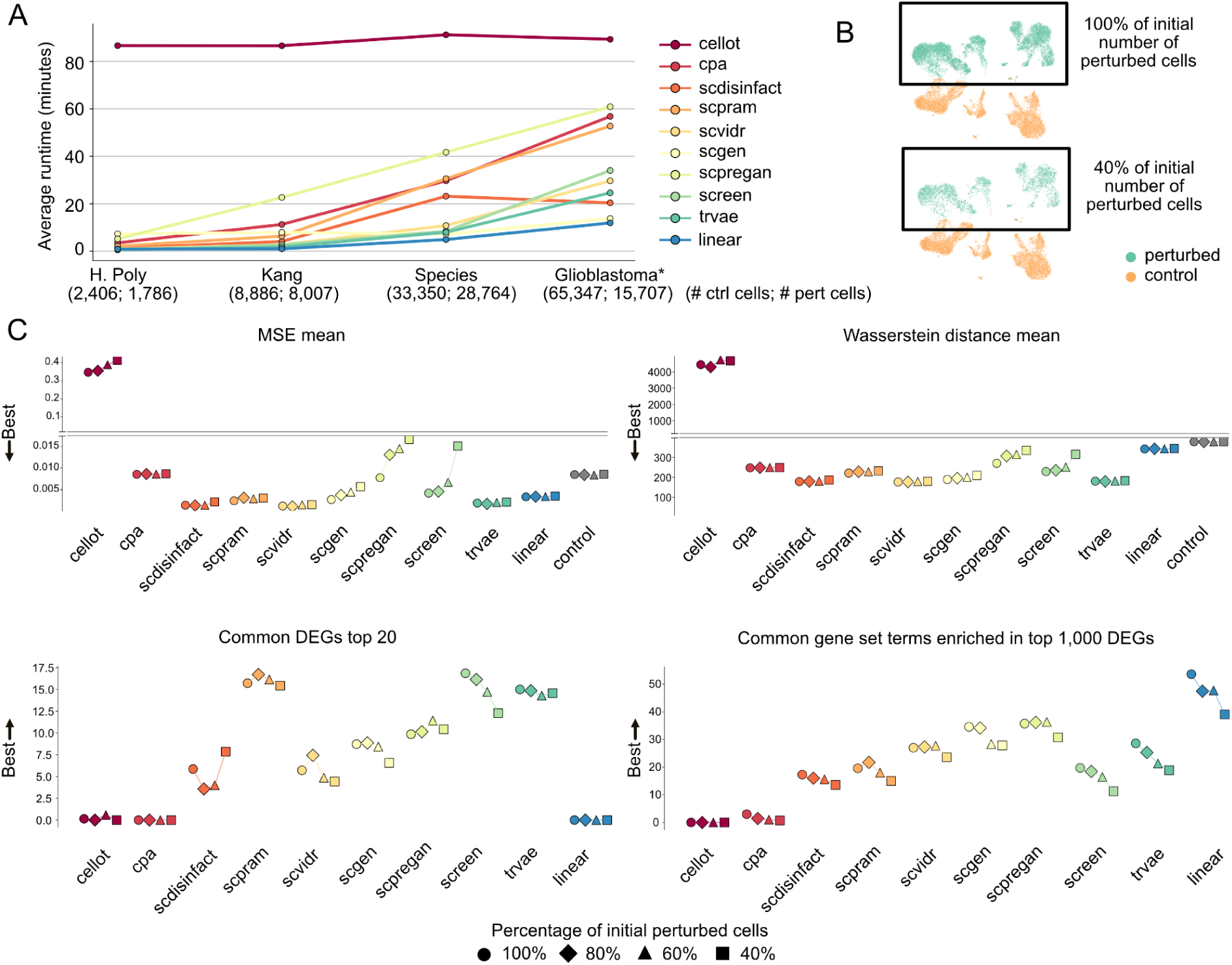
Runtime and ablation experiment performance of tools highlights substantial differences between methods. **A.** Runtime of each tool. *Glioblastoma\** refers to the dataset prior to balancing the number of perturbed and control cells. This distinction is used to illustrate how runtime scales with increasing cell counts. **B.** Ablation experiment design: four datasets were generated from the Kang dataset by progressively reducing the number of initially perturbed cells—100%, 80%, 60%, and 40%. **C.** Performance metrics across these datasets. Top left: Mean Squared Error (MSE). Top right: Wasserstein distance. Bottom left: number of differentially expressed genes (DEGs) shared between predicted and perturbed cells. Bottom right: number of enriched gene sets among the top 1,000 DEGs between predicted and perturbed cells.

We further evaluated model sensitivity to gene set size (Supplementary Fig 10A) by progressively reducing the glioblastoma dataset from all genes to 15,000 and 7,000 highly variable genes (HVGs). As shown in Supplementary Fig 10B, runtimes remained constant. For scVIDR and trVAE, the number of predicted enriched terms decreased with smaller gene sets. In contrast, Wasserstein distance improved for all tools as the gene set was reduced. Meanwhile, the number of DEGs increased, and MSE remained stable, suggesting that errors are largely driven by poorly predicted HVGs, while removing less variable genes has minimal impact on performance.

These results show that while most tools are computationally efficient and robust to moderate reductions in data, performance can vary depending on the nature of the ablation. Notably, reducing input genes or stimulated cells affects biological signal recovery more than standard metrics like MSE, highlighting the importance of evaluating models across multiple dimensions, especially biological relevance.

### Working principle of scArchon to evaluate all the tools on a new dataset

The results above highlight the importance of an open, comprehensive, and fair benchmarking framework that incorporates visual, quantitative, and biological evaluations. As the number of tools for perturbation prediction is constantly growing, we developed scArchon, an accessible platform designed to benchmark models across key performance criteria (https://github.com/hdsu-bioquant/scArchon). Crucially, scArchon eliminates the burden of environment setup by using pre-configured Docker images, enabling seamless integration and execution of existing tools. Running scArchon requires a GPU with CUDA 12.4+ .

Users can select a subset of perturbation prediction tools to benchmark. Given an input dataset, scArchon executes the full pipeline—encompassing model training, prediction, and evaluation. The outputs include: (1) predicted expression matrices in .h5ad format, (2) evaluation metrics in .csv format, and (3) a collection of visualizations, both for individual tools and comparative analyses across models and targets.

The framework is modular and customizable—users can adjust individual tool pipelines, including parameter settings, to better fit their specific experimental design. However, we noted that increasing the number of training epochs generally does not yield significant performance improvements, despite substantially longer runtimes. In line with common practice, most published tools rely on default or fixed parameters across datasets. As such, the parameters used in scArchon serve as a reasonable and fair baseline for initial evaluations.

For more granular or isolated evaluations, users also have the option to run any individual tool independently using its publicly available Docker image.

## Discussion

Out-of-distribution prediction of molecular perturbations represents a significant challenge, given the variety of the possible interventions. It holds the promise to model cell-specific responses to molecular perturbations such as drug treatment or the effect of infection. If successful, this would represent a groundbreaking advance in the field of personalized medicine. It is therefore especially relevant to evaluate to what extent the tools that have been developed in recent years hold this promise (Sharifi-Noghabi et al., 2021). However, evaluating the quality of these predictions represents an equally complex task, which we have tried to address in this work. Previous similar but more limited efforts have been conducted for example as part of the Open Problems in Single-cell Analysis in a collaborative benchmark at NeurIPS 2023. Defining the quality of the predictions is heavily influenced by the choice of qualitative and quantitative evaluation metrics. For example, as we have shown, different dimensionality reduction methods provide sometimes contradicting interpretations. Our findings thus indicate that dimensionality reduction visualisations alone cannot be used as reliable criteria for tool performance evaluation. Therefore, quantitative metrics such as those typically used in model optimization like MSE are needed. According to such metrics, almost half of the tools perform on par or worse than the linear baseline model, in which the perturbation is modelled as a simple constant effect over all cells, highlighting the current limitations in capturing complex perturbation responses. Importantly, several of the metrics which are commonly used are mathematically related and do not represent independent evaluation criteria. Scoring methods based on these metrics might lead to a biased evaluation.

Most metrics take into account more global patterns (e.g. distributional profiles) and might not detect if a tool fails to model relevant biological signals. For this reason, in contrast to previous benchmarks, we also included biological gene-level metrics in our study, such as the number of DEGs or the enriched biological terms amongst those DEGs (L. Li et al., 2024; Wu et al., 2024). Interestingly, evaluating tools based on these more biological criteria sometimes leads to vastly different rankings compared to the typical quantitative metrics, in one way or the other. Some tools perform well on global metrics, but detect enriched biological terms that are not related to the underlying biology. The correlation between the rankings of the quantitative metrics and the biological metrics is low, emphasizing this discrepancy (< 0.5, Fig 3C). We believe that this is similar to the hallucination effects that are common in complex deep learning approaches such as LLMs and computer vision (Liu et al., 2024). Unlike hallucinations in LLMs which generate plausible but false text, biological hallucinations in perturbation models manifest as spurious pathway enrichments that could mislead downstream biological interpretation and experimental design. Including biological priors into the prediction models might alleviate these effects, as they would constrain the predictions to biological plausible solutions (Doncevic & Herrmann, 2023; Ma et al., 2018; Seninge et al., 2021). These observations underscore the necessity for novel evaluation approaches that better represent the capacity of perturbation prediction tools to model biological responses.

As expected, the performance of each tool appears to be highly dependent on the type and strength of the perturbations, varying considerably across different testing scenarios. To address this, we conducted evaluations using a broad range of data types and experimental conditions. This approach aims to assess not only the overall performance but also the generalizability of each tool. Such diversity in testing is crucial, as many tools demonstrate high accuracy in tasks in which the effect to be predicted is the main source of variance yet struggle when faced with scenarios in which this effect is masked by other confounding effects, like patient specific responses. In this benchmark, we chose to focus on molecular perturbations that are distinct from genetic perturbations such as gene knock-outs obtained from CRISPR-screens, which have been evaluated elsewhere (Ahlmann-Eltze et al., 2024). We believe that drug perturbation prediction is a more challenging task, as the mode of action of drug perturbations is less clear than when well defined genes are altered.

Overall, taking into account the different types of metrics introduced here, we observe that the tools that consistently performed best in our benchmarking study are scVIDR, scPRAM, scGen, scDisInFact, and trVAE.

Given the limited overall performance of the tools, especially in more challenging datasets, predicting the full transcriptional response of each cell might represent an overly ambitious and sometimes unnecessary objective. Alternatively, predicting the changes in the activities of specific pathways of interest or the drug sensitivity of cells appears as a more realistic goal which could be sufficient in many application scenarios (C. Li et al., 2024; Burkhardt et al., 2021; Fustero-Torre et al., 2021; Hsieh et al., 2023).

As the number of tools to predict perturbations in single-cell datasets will continue to increase, we are proposing the scArchon framework, built on Snakemake (Mölder et al., 2021), which enables efficient and reproducible testing of perturbation prediction tools across diverse datasets, based on multiple metrics. Its modular design allows for seamless integration of new tools and datasets, promoting large-scale, standardized evaluations. Such a common benchmarking system would not only facilitate transparency in reporting but also accelerate methodological improvements by providing a clear reference for performance assessment (Mahmood, 2025). scArchon contributes to this vision by offering a unified platform that enables systematic, reproducible, and transparent evaluation of perturbation prediction tools. As the field progresses, the availability of such a framework could serve as a foundation for measuring improvements in prediction accuracy and biological relevance, fostering innovation and guiding the development of more robust models. Therefore, we encourage the authors of future methods to make use of this unbiased evaluation platform and provide dockerized images of their tools.

## Material and Methods

### Datasets

#### Kang dataset

The Kang dataset (Supplementary Fig 1) originates from human peripheral blood mononuclear cells (PBMCs) comprising seven different immune cell types and is publicly available through the GEO repository under accession number GSE96583 (Kang et al., 2017). For our work, we used the pre-processed and annotated version provided by Lotfollahi et al. (Lotfollahi et al., 2019), which can be downloaded from https://www.dropbox.com/s/wk5zewf2g1oat69/train_pbmc.h5ad?dl=1. The dataset includes 16,893 cells in total, 8,007 control cells and 8,886 interferon-β perturbed cells, and covers 6,998 genes.

#### H. Poly dataset

The H. Poly dataset (Haber et al., 2017) (Supplementary Fig 2) consists of intestinal epithelial cells from mice that were infected with the parasitic helminth *Heligmosomoides polygyrus* over a ten-day period. The dataset is available as raw data under GEO accession number GSE92332. We used the processed and annotated version published by Lotfollahi et al. (Lotfollahi et al., 2019), which can be accessed at https://www.dropbox.com/s/7ngt0hv21hl2exn/train_hpoly.h5ad?dl=1. We retained only cell types with more than 400 cells to ensure statistical robustness and avoid biases due to small sample sizes. Thus, the final dataset contains 2,406 control cells and 1,786 perturbed cells from five different cell types, and 7,000 genes.

#### Species dataset

The species dataset (Hagai et al., 2018) (Supplementary Fig 3) (accession id E-MTAB-6754) consists of cells from the four species mouse, rabbit, pig, and rat treated with lipopolysaccharide (LPS) for six hours. The processed data was downloaded from https://www.dropbox.com/s/eprgwhd98c9quiq/train_species.h5ad?dl=1 which was published by (Lotfollahi et al., 2019). The dataset consists of 62,114 cells (33,350 control and 28,764 perturbed) and 6,619 genes.

#### Glioblastoma dataset

The Glioblastoma dataset (Supplementary Fig 4) contains single-cell transcriptomic profiles of glioblastoma patient-derived cells and is available as raw data under GEO accession number GSE148842. We used the annotated version by Levitin et al. (Levitin et al., 2023), which can be downloaded from https://drive.google.com/file/d/18-KInmm43wKdBX95Gq9xbuzAQwtLjgE9/view?usp=sharing. In our analysis, we subset the dataset to include only five patients (PW030, PW032, PW034, PW036, and PW040) that had both control and panobinostat-treated conditions available, excluding all other treatment groups. The original dataset consists of 81,054 cells, of which 65,347 are control and 15,707 are perturbed, and 20,098 genes. However, we downsampled the control cells of the dataset to match the number of perturbed cells.

### Dataset preprocessing

While some studies propose evaluating model performance on a subset of genes, such as the most highly variable or the most differentially expressed ones (L. Li et al., 2024), we chose to conduct our benchmark across the full set of genes. We reasoned that restricting the analysis to selected genes may emphasize biologically prominent features, but it can also introduce bias and misrepresent the model’s generalizability. All datasets were count normalized and log-transformed.

### Model selection

#### Benchmarked tools

Each of the selected methods employs deep learning and relies on dimensionality reduction to project high-dimensional gene expression data into a latent representation where it then learns a mapping from control to perturbed state.

scGen (https://github.com/theislab/scgen, commit d79e1f0, version 2.1.0) uses a variational autoencoder (VAE) framework in which the perturbation effect is modeled as a simple linear vector shift in the latent space that can then be applied to new control samples to simulate the perturbed state.

CellOT (https://github.com/bunnech/cellot, commit 522d2b9), scPRAM (https://github.com/jiang-q19/scPRAM, commit a462429), scVIDR (https://github.com/BhattacharyaLab/scVIDR, commit 605825f), and SCREEN (https://github.com/Califorya/SCREEN, commit c0dc236) use optimal transport (OT) in the latent space to learn a transport map that shifts the distribution of control cells toward that of perturbed cells.

CPA (Compositional Perturbation Autoencoder) (https://github.com/theislab/cpa, commit fbd7c02) disentangles confounding variables—such as cell type, patient, or batch—from the perturbation effect in the latent space by separately learning them through embedding layers. It makes the assumption that the latent space can then be described as the sum of the different components.

scDisInFact (https://github.com/ZhangLabGT/scDisInFact, commit 9466ecf) also learns a disentangled latent space that separates batch effects from condition-specific signals. It addresses three tasks simultaneously: batch effect removal, identification of condition-associated key genes, and perturbation prediction. This design makes scDisInFact applicable to studies with multiple batches and experimental conditions, where integrated analysis is required.

trVAE (transfer variational autoencoder) (https://github.com/theislab/trVAE_reproducibility, commit `2a594b6`) extends the CVAE architecture by modeling the perturbation effect as a condition vector, which is disentangled from batch variables such as cell state, donor, or species. These vectors are integrated in the decoder and optimized using a maximum mean discrepancy (MMD) loss, which aligns the latent representations across different conditions.

scPreGAN (https://github.com/XiajieWei/scPreGAN, commit d9fa8ea) employs a generative adversarial network (GAN) architecture, combining it with an encoder to learn a generative mapping from control to perturbed states. The GAN framework encourages the model to produce perturbed cell states that are both realistic and aligned with the underlying biological signal, using adversarial training to improve fidelity.

Tools were used as originally described in their respective publication using their default parameters. For scGen we removed early stopping. For CPA, we modified the model following guidance found in the CPA GitHub repository (Supplementary Notes).

### Linear model baseline

For each gene, we compute the average expression across all cells in the control and perturbed conditions within the training set. The difference between these two averages represents the estimated perturbation effect for that gene. To generate predictions on the test set, we add this gene-wise difference to the expression profile of each test cell from the control condition.

### Control baseline

We used the control condition directly by applying the same metrics to the unmodified control cells compared to the true perturbed cells. For metrics that rely on differential comparisons, such as those requiring DEG analysis between control and perturbed states, scores for the control baseline could not be computed

### Evaluation metrics

#### Dimensionality reduction techniques

Principal component analysis was performed using the function <u>scanpy.pp</u>.pca (Wolf et al., 2018) with the number of components set to 50 and random_state set to 42. UMAP was computed using scanpy.tl.umap with min_dist set to 0.5, spread set to 1, and random_state set to 42. t-SNE was applied using scanpy.tl.tsne with random_state set to 42.

### Metrics

MSE, MMD, Wasserstein distance, t-test score, pearson distance and cosine distance were computed using the pertpy.tl.Distance function from pertpy (Heumos et al., 2024). R² was computed using the implementation from scButterfly (Cao et al., 2024) (https://github.com/BioX-NKU/scButterfly, commit eb31e04). We calculated R² across three gene subsets: the top 20 DEGs, the top 100 DEGs, and the entire set of genes.

### Ranking methodology and tool score

The tools were ranked based on the average ranks across five metrics: MSE, Wasserstein distance, t-test, common DEGs and common enriched terms. Since the average rank was not intuitive to rank the different tools as the smallest value was representing the best tool, we reversed the score by using (1-(avg_rank-1))/(number_of_tools-1) so as to keep a score between 0 and 1, where the greatest value is for the tool which scored best overall.

### Biological evaluation

Enrichment analysis was performed on the top 1000 DEGs identified using the scanpy.tl.rank_gene_groups function with the method parameter set to t-test. Gene set enrichment analysis was conducted using the gseapy package (Fang et al., 2022). Gene sets were taken from the MSigDB resource (Liberzon et al., 2011), specifically the MSigDB_Hallmark_2020 collection, and from Gene Ontology (Ashburner et al., 2000), including GO_Biological_Process_2021, GO_Molecular_Function_2021, and GO_Cellular_Component_2021. For the H. Poly dataset, organism variable was set to Mouse. Enrichment terms were retained if the adjusted p-value was below 0.05. For each significant term, gene scores were calculated using scanpy.tl.score_genes, which computes the average expression of the selected gene set subtracted by the average expression of all other genes.

### Construction of scArchon

scArchon was implemented in Snakemake (v8.28.0) and utilizes containerized environments available at https://hub.docker.com/u/hdsu. These Docker containers include all necessary software dependencies for the supported tools. The tools require CUDA versions ranging from 11.6 to 12.4; therefore, systems equipped with CUDA 12.4 or later are compatible with all components.

Execution of the pipeline also requires Singularity (Kurtzer et al., 2017), tested on versions 3.6 and 4.1, with newer versions expected to be compatible. On average, each image occupies approximately 6.5 GB of disk space. Running for example four tools thus requires a minimum of ∼30 GB to accommodate both images and output files. Once downloaded, the Singularity images are cached locally and reused, so subsequent runs do not incur additional download time.

Tool runtimes vary considerably. scPreGAN, scDisInFact, scPRAM, and scVIDR execute relatively quickly. CellOT is significantly slower due to the absence of GPU support and the fact that scGen has to be trained prior to extracting the latent space representation of the dataset. SCREEN and CPA also require extended runtime. Based on empirical results, it is recommended to exclude CellOT, SCREEN, and CPA from initial evaluations.

## Code Availability

scArchon’s implementation can be found at https://github.com/hdsu-bioquant/scArchon.

## Author contribution

JR and RD designed the study. JR developed the scArchon benchmarking platform. JR and RD implemented the snakemake pipeline. JR and RD carried out the analyses. RD, DD, AL and CH provided guidance and feedback throughout the study. DTB tested scArchon. All authors contributed to the writing of the manuscript.

## Acknowledgements

JR is supported by the Carl-Zeiss-Stiftung through the AI-CARE project (P2022-08-008). RD is supported by the Deutsche Forschungsgemeinschaft (DFG) through the special priority program SPP2202. AL is funded through a PhD grant from the Studienstiftung des Deutschen Volkes.

