## Supplementary Figs for "Tracking biological hallucinations in single-cell perturbation predictions using scArchon, a comprehensive benchmarking platform"

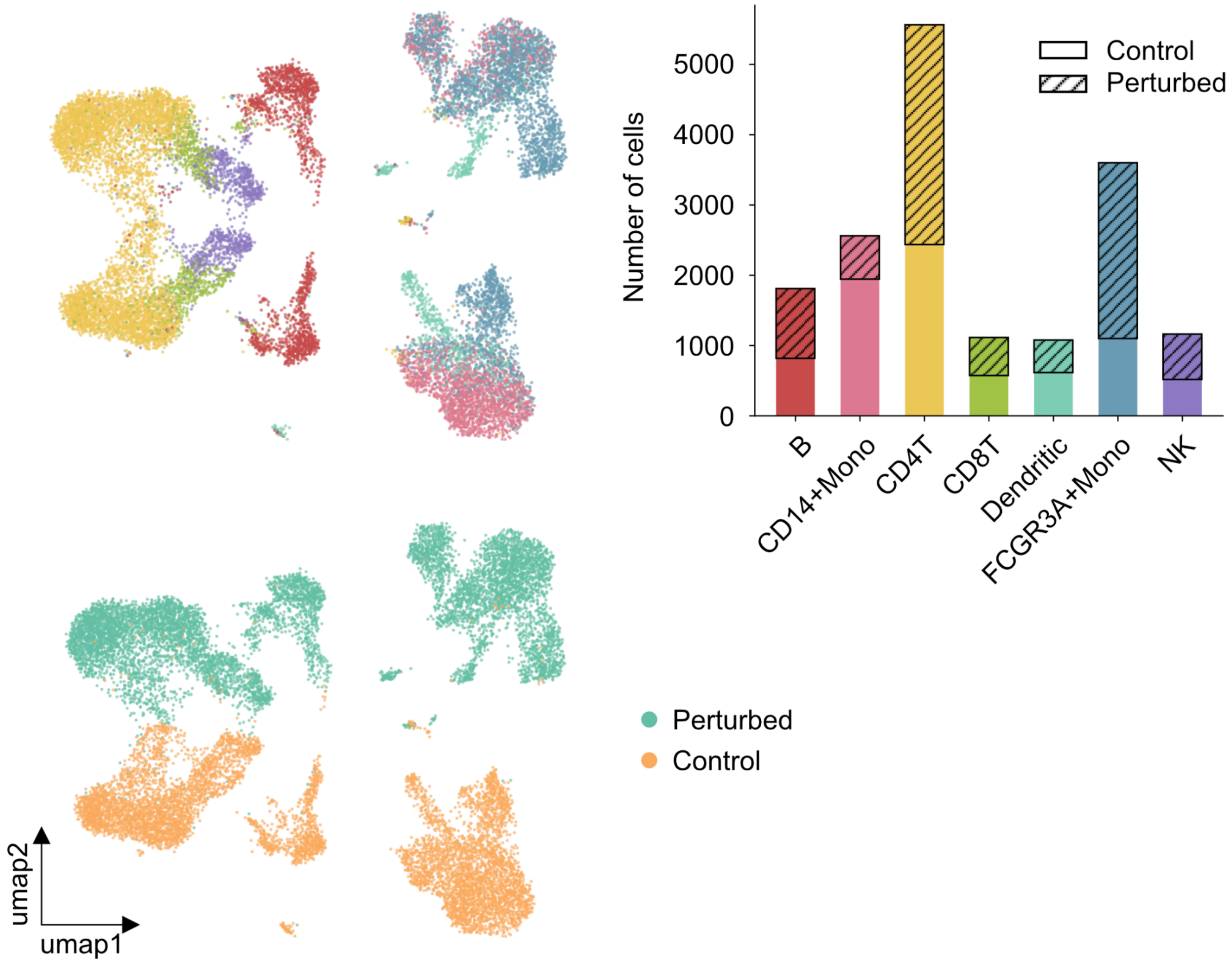


**Fig S1:** Overview of the Kang dataset.
The top-left panel displays the full dataset, colored by cell type. The corresponding cell type distribution is summarized in the top-right bar plot. The bottom-left panel highlights the distribution of cells across the perturbed and control conditions. The dataset includes 16,893 cells, from which 8,886 cells in control and 8,007 cells in perturbed, and 6,998 genes.

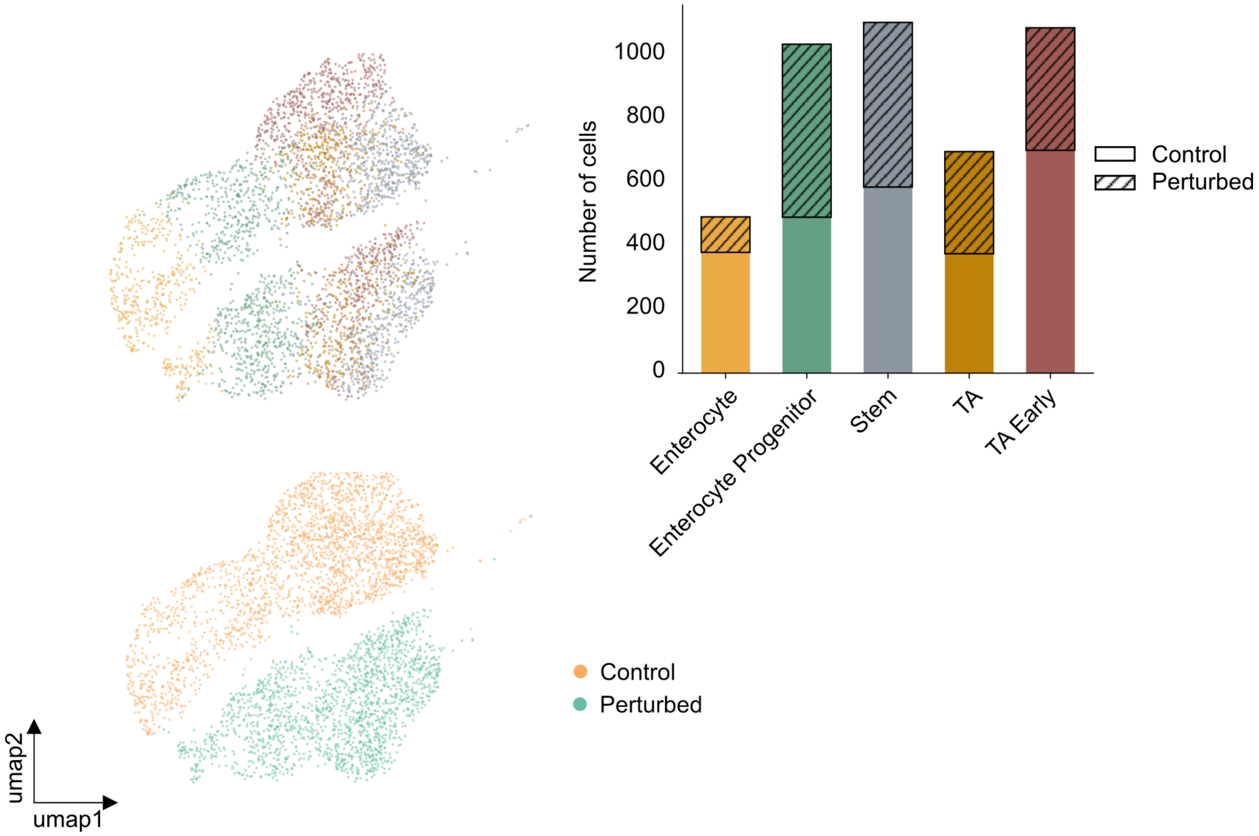


**Fig S2:** Overview of the H. Poly dataset.
The top-left panel displays different cell types, colored according to the distribution shown in the top-right bar plot. The bottom-left panel illustrates the separation between control and perturbed cells. The dataset contains 4,192 cells and 7,000 genes, including 2,406 control cells and 1,786 perturbed cells.

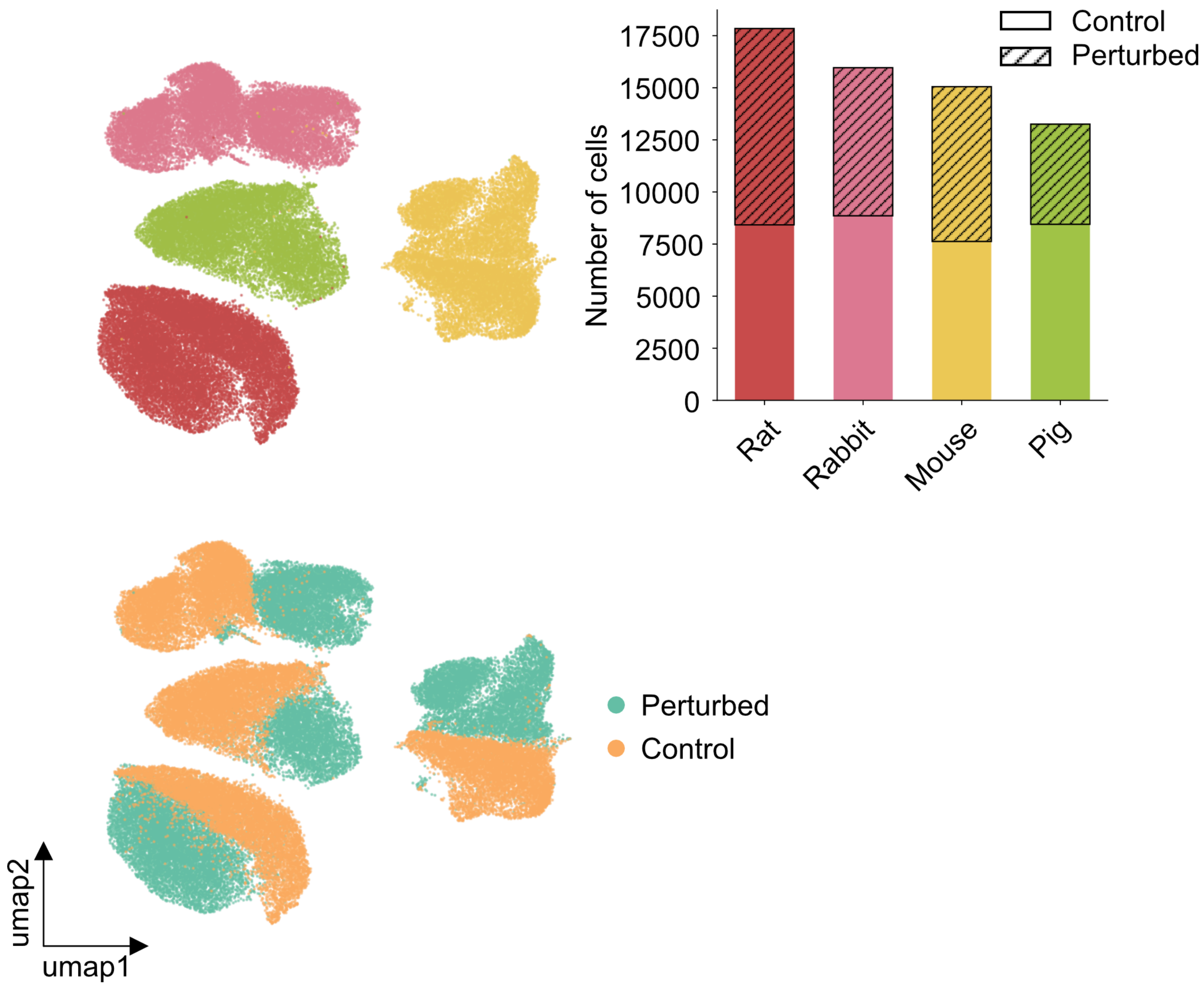


**Fig S3:** Overview of the Species dataset.
The top-left panel displays different cell types, colored according to the distribution shown in the top-right bar plot. The bottom-left panel illustrates the separation between control and perturbed cells. The dataset consists of 62,114 cells and 6,619 genes, including 33,350 control cells and 28,764 perturbed cells.

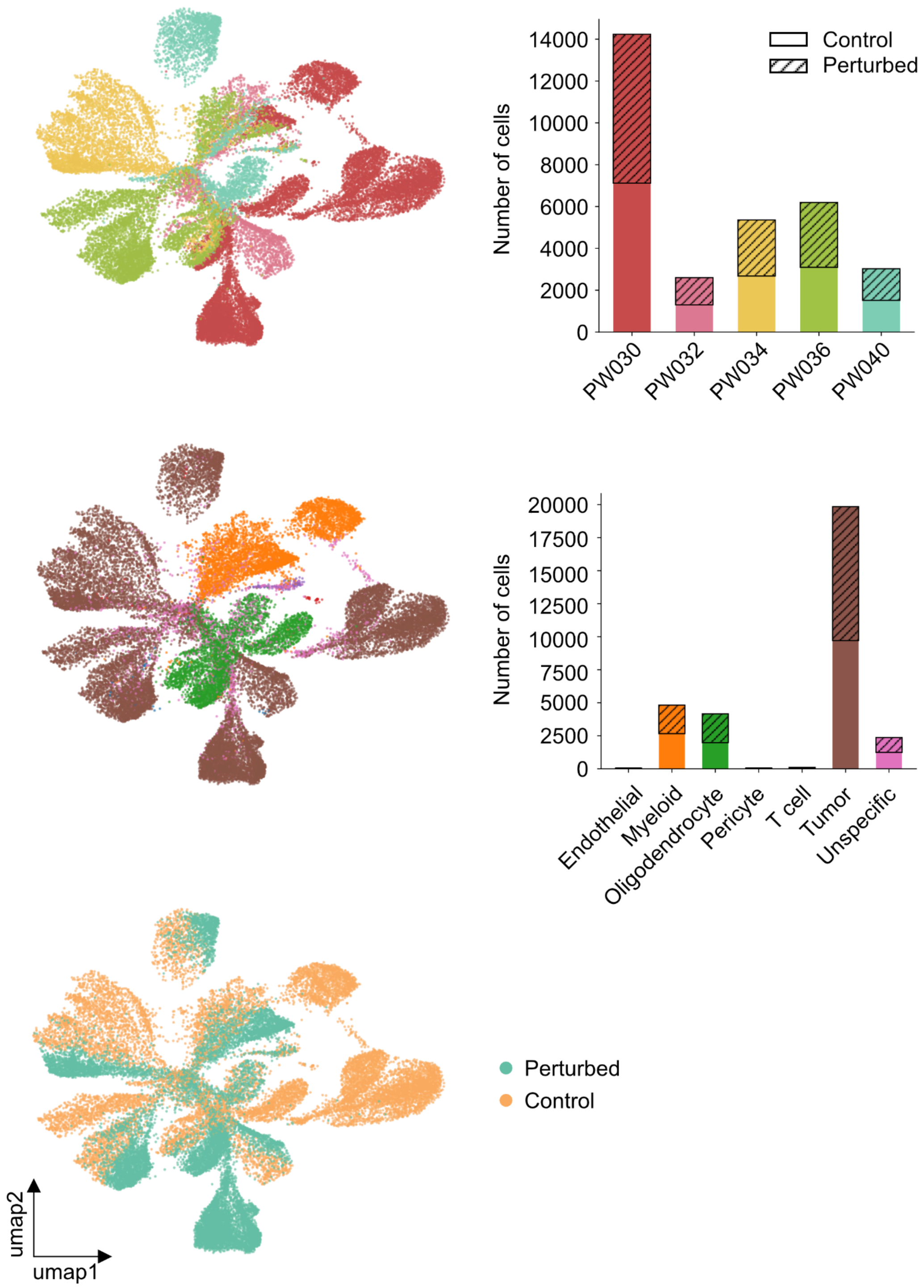


**Fig S4:** Overview of the Glioblastoma dataset.
The top-left panel shows different cell types, colored as in the bar plot on the top-right. The bottom-left panel displays control and perturbed cells. The dataset comprises 31,414 cells and 7,000 genes, with patient and cell type annotations. It includes 15,707 control cells and 15,707 perturbed cells.

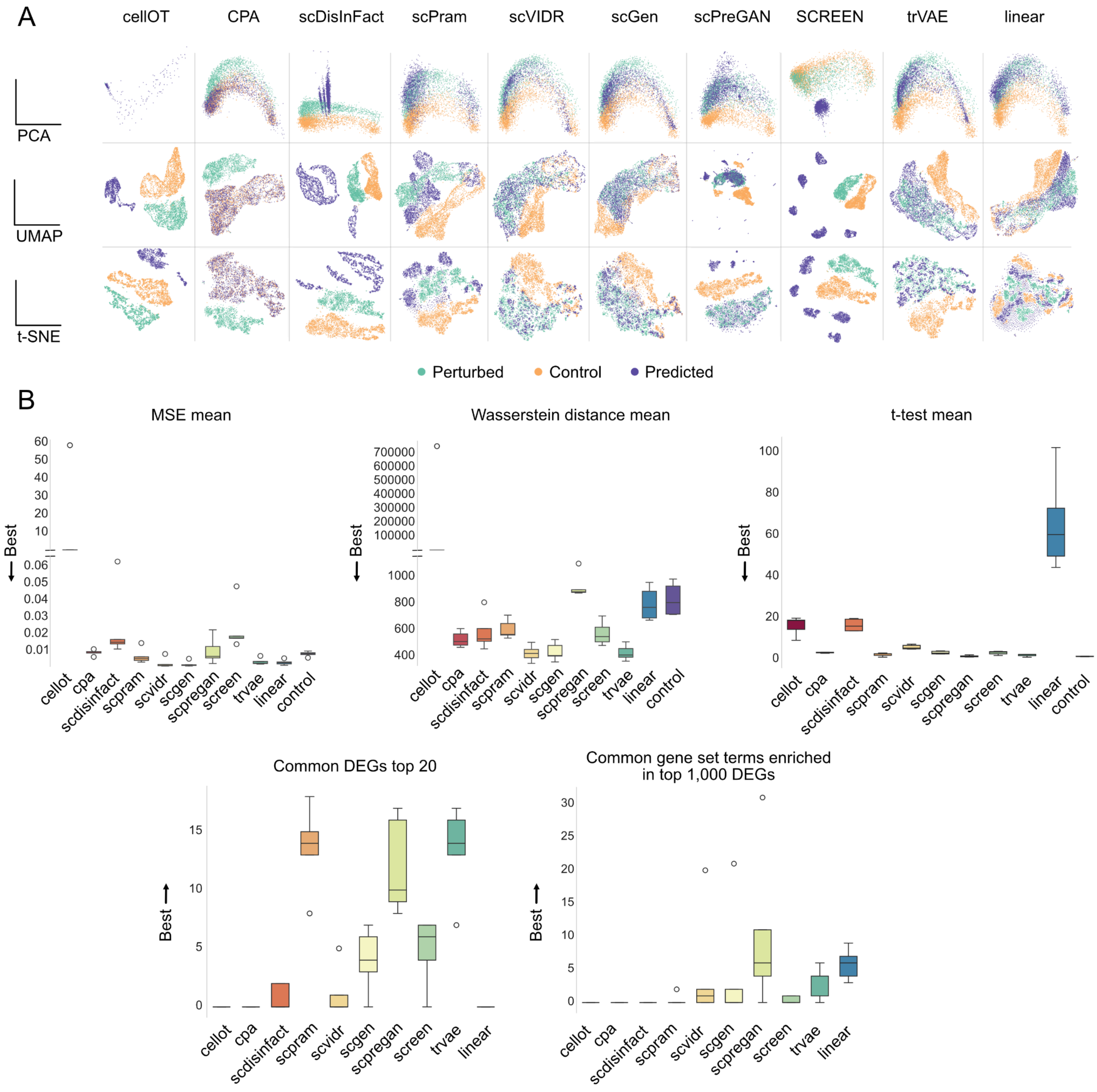


**Fig S5:** Results on the H. Poly dataset.

**A.** Dimensionality reduction plot showing the aggregated outputs from all experiments conducted on the dataset.

**B.** Evaluation metrics including MSE, Wasserstein distance, t-test, overlap of top 20 differentially expressed genes (DEGs), and overlap of top 1,000 DEGs used for enrichment analysis.

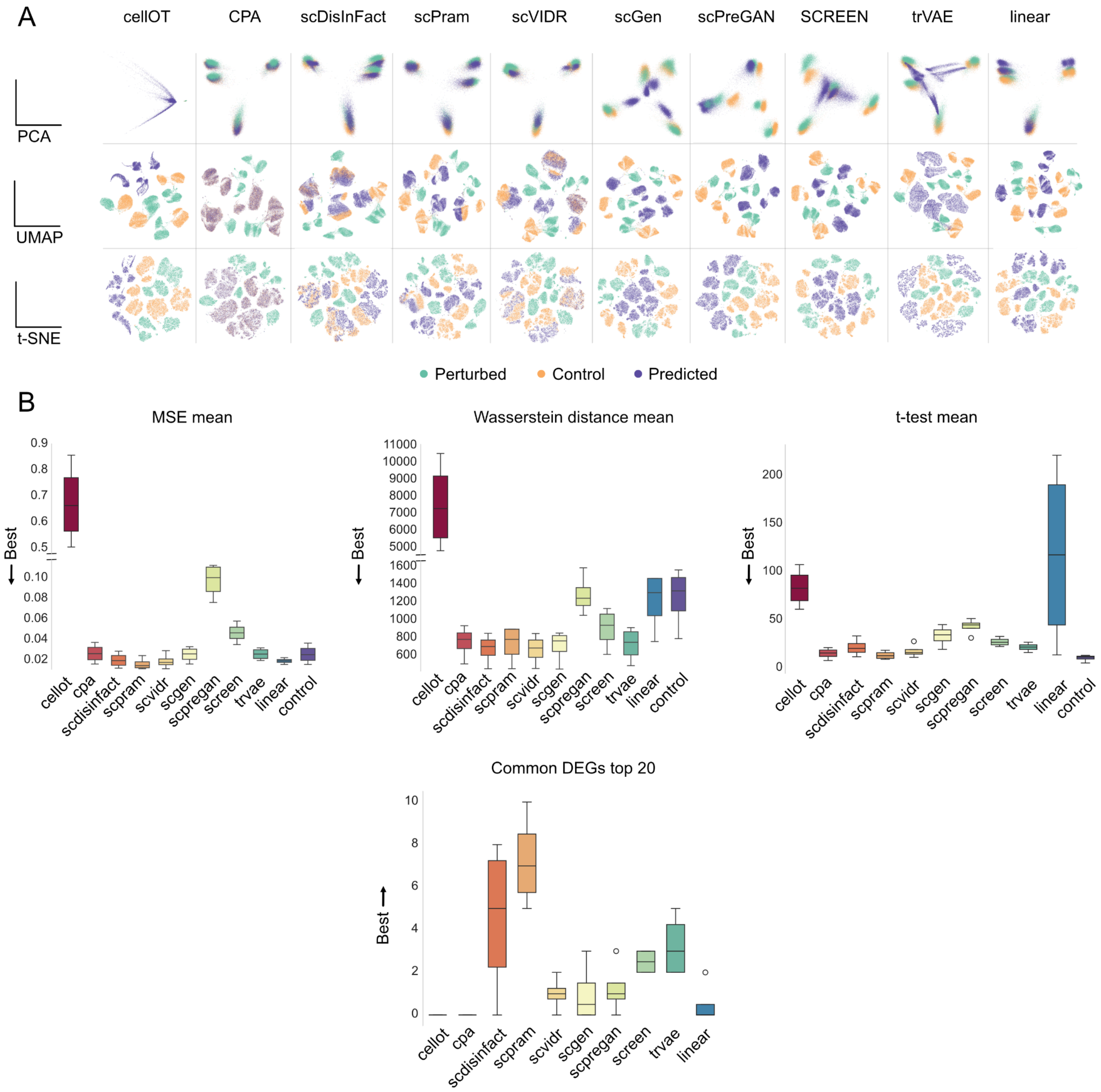


**Fig S6:** Results on the Species dataset.
**A.** Dimensionality reduction plot showing the aggregated outputs from all experiments conducted on the dataset.
**B.** Evaluation metrics including MSE, Wasserstein distance, t-test and overlap of top 20 differentially expressed genes (DEGs).

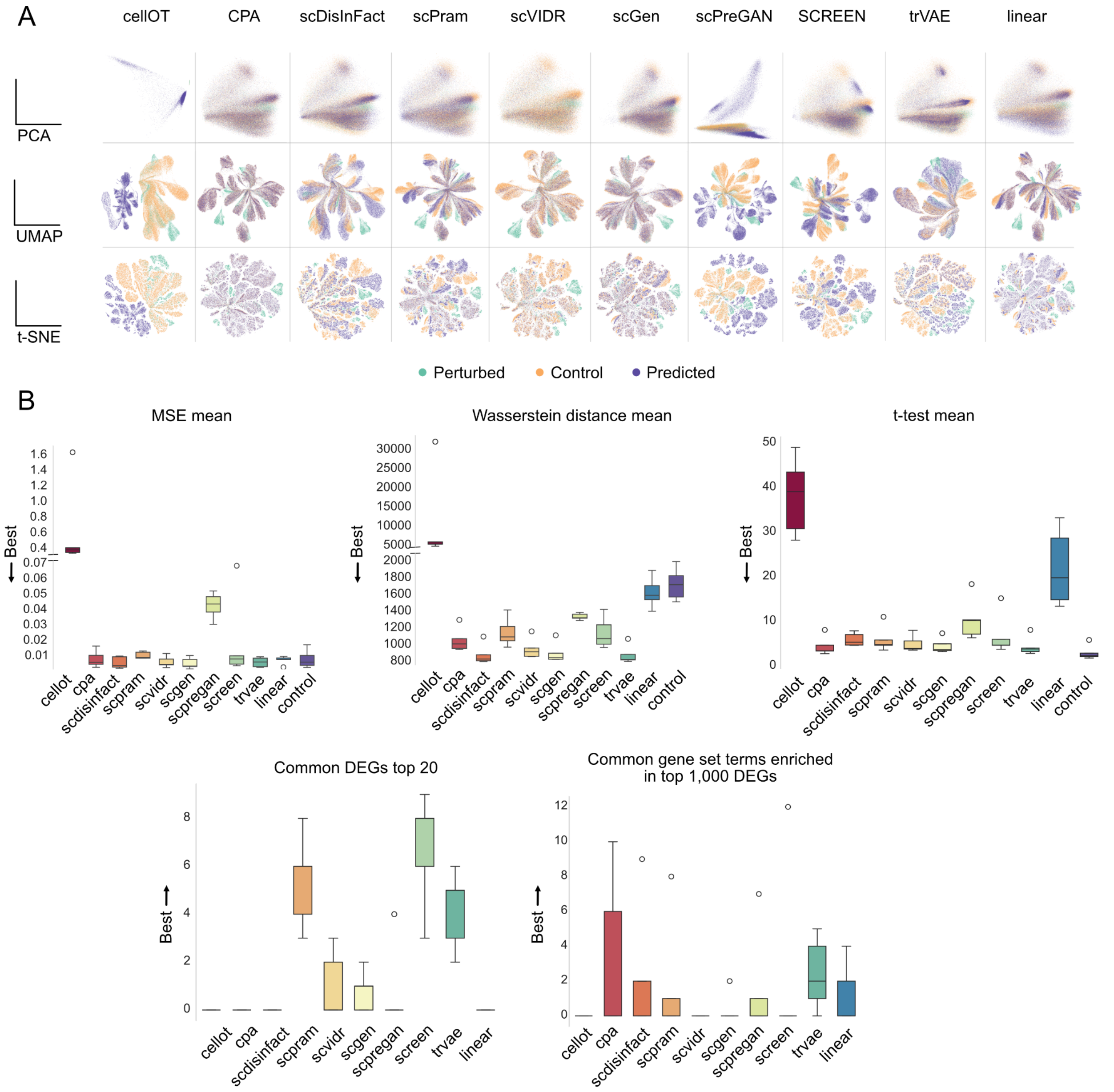


**Fig S7:** Results on the Glioblastoma dataset.

**A.** Dimensionality reduction plot showing the aggregated outputs from all experiments conducted on the dataset.

**B.** Evaluation metrics including MSE, Wasserstein distance, t-test, overlap of top 20 differentially expressed genes (DEGs), and overlap of top 1,000 DEGs used for enrichment analysis.

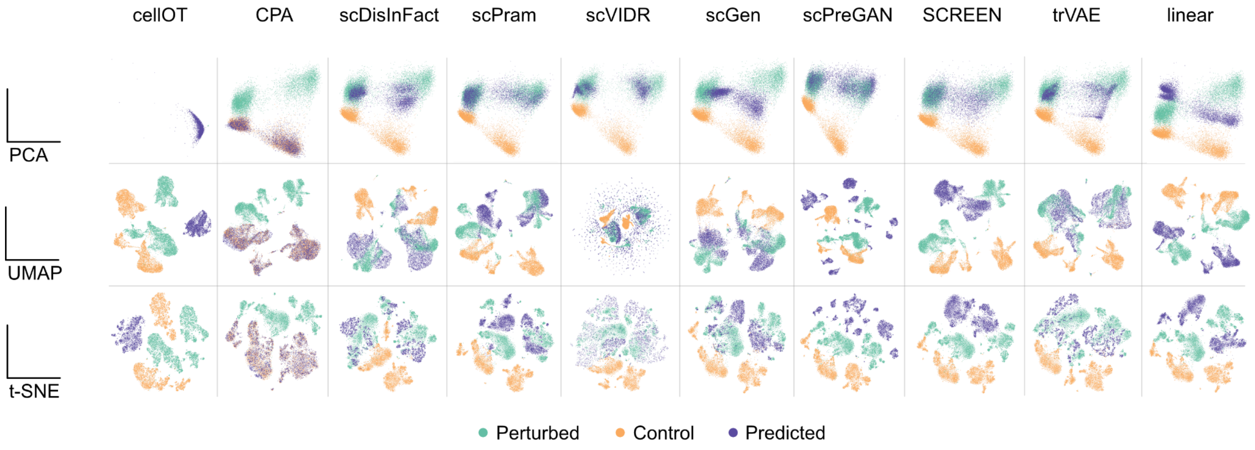


**Fig S8:** Dimensionality reduction plot showing the aggregated outputs from all experiments conducted on the Kang dataset.

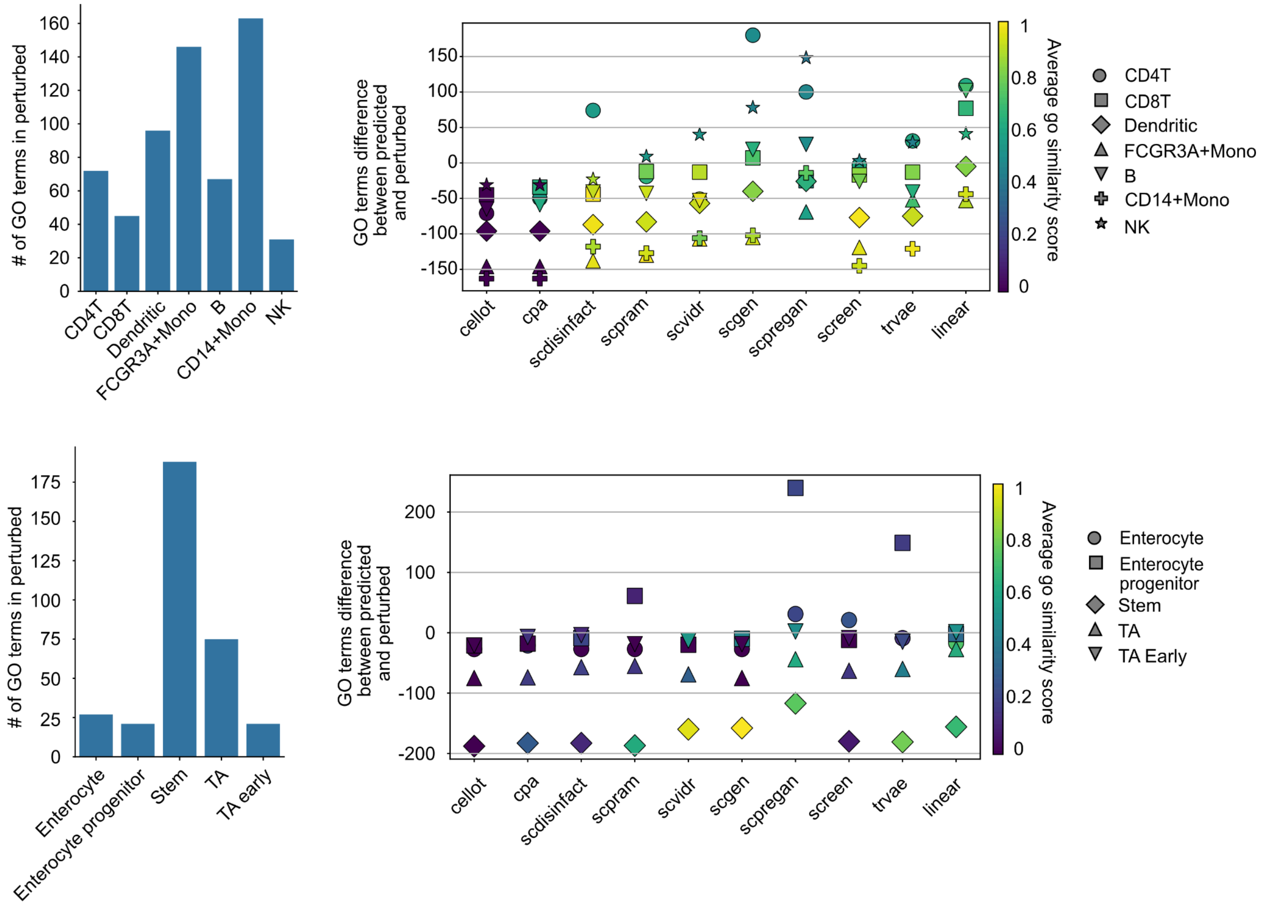


**Fig S9:** Similarity scores for the two datasets with enriched GO terms. The top panel shows scores for the Kang dataset and the bottom panel for the H. Poly dataset.

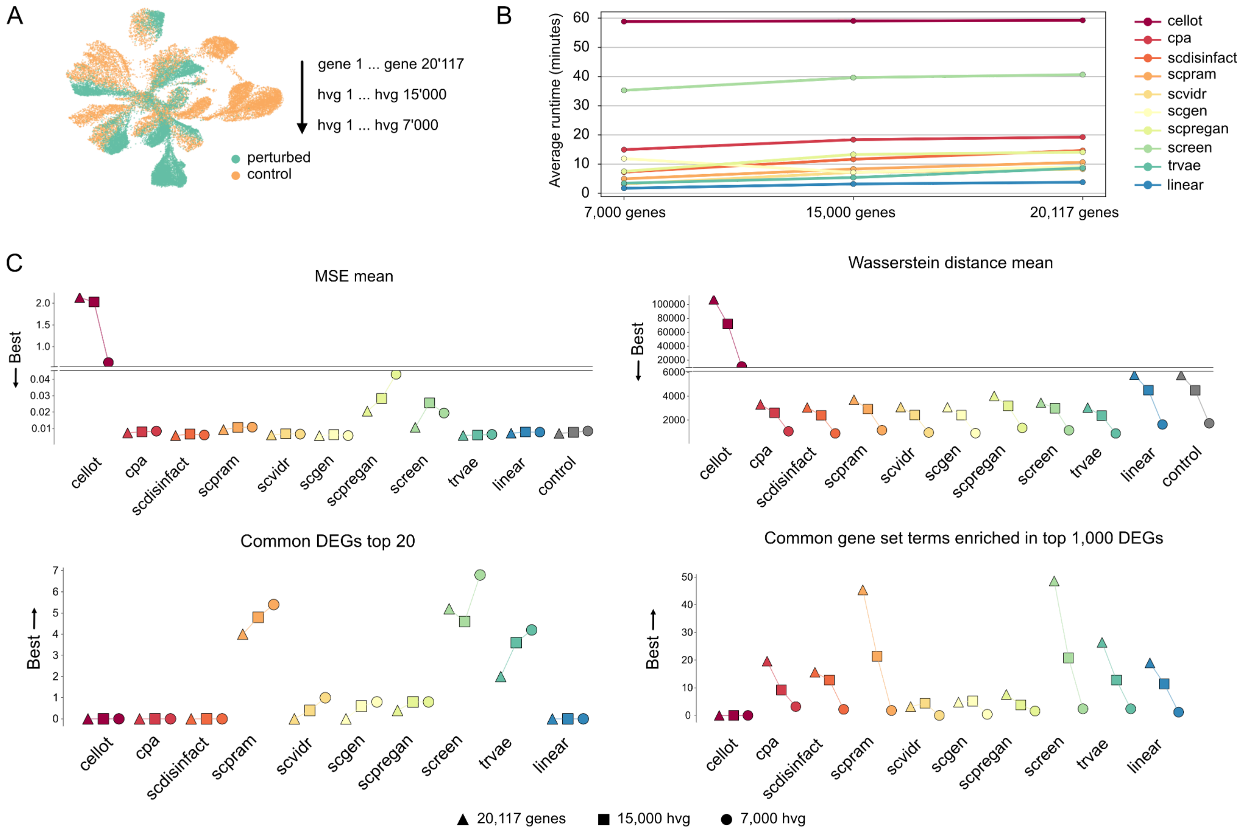


**Fig S10: Gene ablation experiment.**

**A.** Schematic of the experiment. Starting with the Glioblastoma dataset, we progressively reduce the number of input genes—from the full set to the top 15,000 highly variable genes (HVGs), and then to the top 7,000 HVGs.

**B.** Runtime comparison of the different tools across datasets with varying gene counts.

**C.** Metric trends as gene count decreases. Top left: Mean Squared Error (MSE); top right: Wasserstein distance; bottom left: number of differentially expressed genes (DEGs) shared between prediction and reference; bottom right: number of shared enriched gene sets.
