## Supplementary Notes for "Tracking biological hallucinations in single-cell perturbation predictions using scArchon, a comprehensive benchmarking platform"

### Reproduction

Before conducting tool comparisons, we first sought to reproduce key results from each method's original publication. When available, we replicated either the published vignette or a representative Fig from the paper to serve as a sanity check, ensuring that each tool was implemented and executed correctly. Any deviations from the original code were documented and justified accordingly.

#### scGen reproduction

By successfully reproducing their vignette (Fig Supplementary Notes (SN) 1), we confirm that our pipeline produces consistent outputs with the original implementation. The vignette can be found at: https://scgen.readthedocs.io/en/stable/tutorials/scgen_perturbation_prediction.html. As stated in Supplementary Table 1 of the scGen publication, the same model architecture is applied across all experiments. Since scGen does not require dataset-specific fine-tuning, we used the default parameters for this analysis.

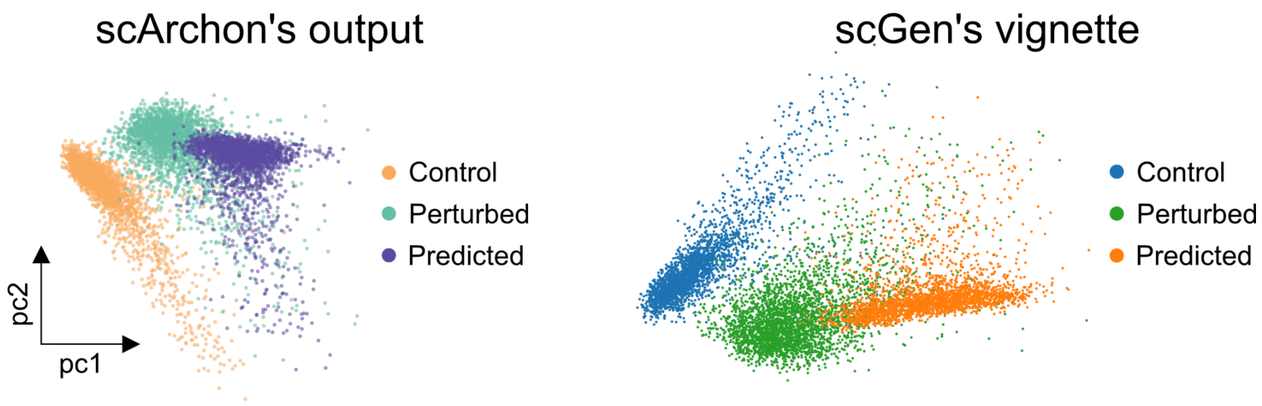


**Fig SN 1:** Comparison of scGen results on the CD4T cell type from the Kang dataset.
The left panel shows the output obtained using our pipeline, while the right panel presents the result from the official scGen vignette.

### trVAE reproduction

The original trVAE vignette, available at github/theislab/trvaep/blob/master/example/sample_notebook.ipynb, relies on an older version of the dataset that is incompatible with current versions of Scanpy. To reproduce the results, we used the version of the Kang dataset from our study and adapted the vignette accordingly. Notably, the original vignette predicts using the *perturbed* data as input, which is biologically incorrect. We corrected this by modifying the predict function to use *control* data as input instead. This proper implementation is found on the official trVAE reproducibility scripts https://github.com/theislab/trVAE_reproducibility/blob/master/scripts/train_trVAE.py, where the perturbed condition is excluded from training and used for prediction. While trVAE's original setup uses the top 3,000 highly variable genes, we removed this filtering step to maintain consistency with scArchon's input data. The resulting output aligns with the original (Fig SN2), though slight differences in representation appear due to variations in the plotting functions used.

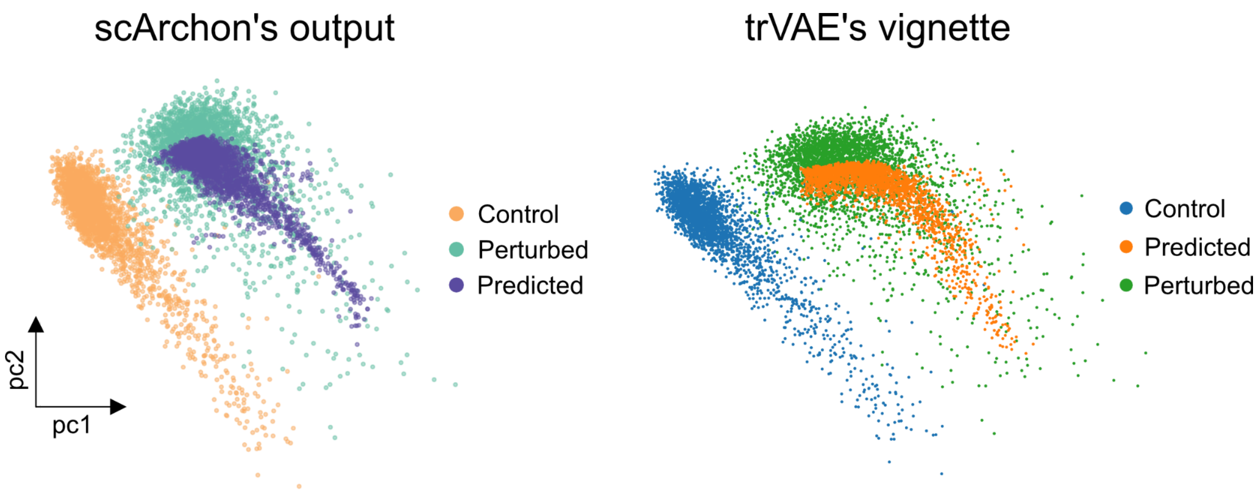


**Fig SN 2:** Reproduction of trVAE results on the Kang dataset. Cell type: CD4T.

### scPreGAN reproduction

We reproduce results from the reproducibility code from scPreGAN adapted from https://github.com/XiajieWei/scPreGAN-reproducibility/scPreGAN_OOD_prediction.py. We show that the implementation found in scArchon matches the results obtained from the reproduction script (Fig SN3).

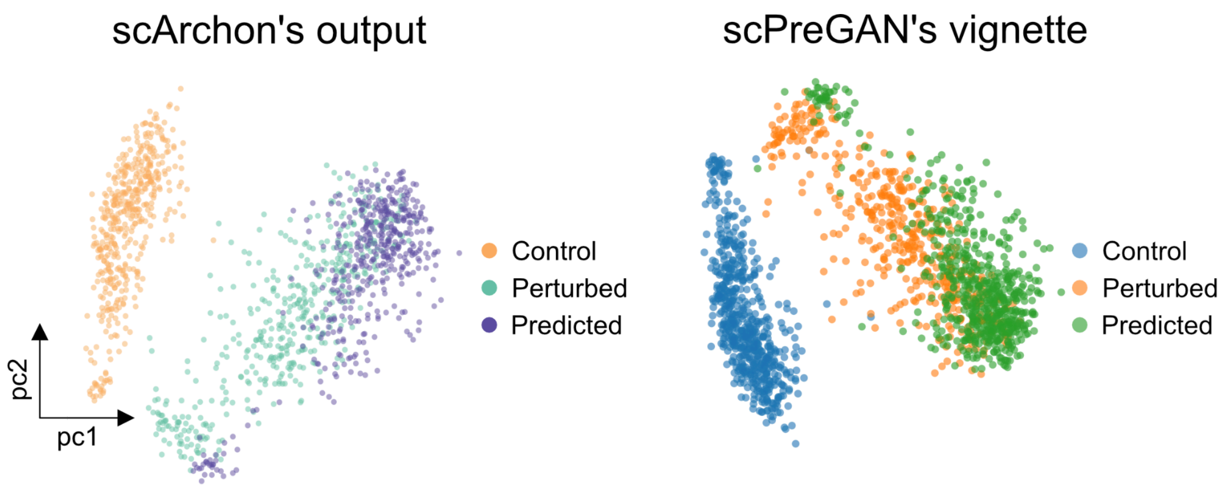


**Fig SN 3:** Reproduction of scPreGAN results on the Dendritic cell type from the Kang dataset.

**scVIDR reproduction**

The scVIDR vignette, available at https://github.com/BhattacharyaLab/scVIDR/blob/main/notebooks/SupplementalFig3.ipynb, was run on Kang B cells. On Fig SN4, the left panel displays the result obtained using the scArchon pipeline, while the right panel shows the output from the original scVIDR vignette.

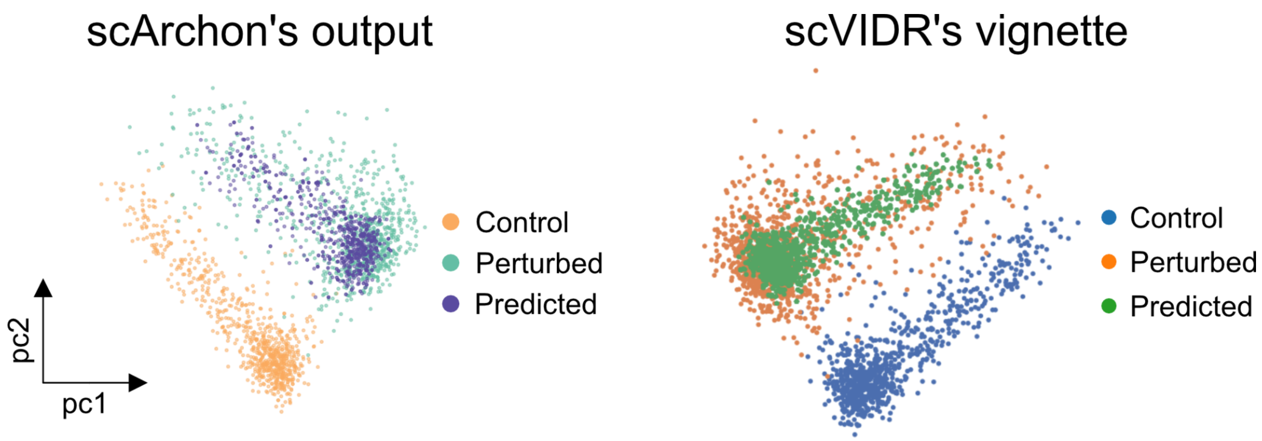


**Fig SN 4:** Reproduction of scVIDR results on B cells from the Kang dataset.

### CellOT Reproduction

In Fig SN5, the left panel reproduces results from cellOT's original publication (Fig 2 in cellOT's paper), while the right panel presents outputs generated using the scArchon pipeline. We show both the original implementation of cellOT (which does not incorporate control data) and an alternative version that includes control data. Results are presented for two conditions: *imatinib* (top row) and *trametinib* (bottom row). It is worth noting that the 4i dataset used in this benchmark contains only 48 features—substantially fewer than the thousands of genes typically present in single-cell RNA sequencing (scRNA-seq) datasets.

Our reproduction confirms that scArchon successfully replicates the results reported in cellOT's original study on the 4i dataset, showing a strong alignment between predicted and true perturbed states. Given this performance, one might expect cellOT to perform similarly well on more complex, higher-dimensional scRNA-seq data—especially when dimensionality is reduced using a learned representation such as that provided by scGen. However, in line with findings from previous benchmarks found in scPRAM' (Fig 2C in scPRAM's paper) and scVIDR's (Fig 2B in scVIDR's paper) papers, our results indicate that cellOT exhibits reduced performance on scRNA-seq datasets, suggesting that its strong results on lower-dimensional data may not generalize to more complex transcriptomic profiles. Additionally, we note that cellOT's usability remains a challenge, as highlighted in a recent preprint (Li et al., 2024), which states implementation difficulties as the reason for excluding the tool from their benchmark.

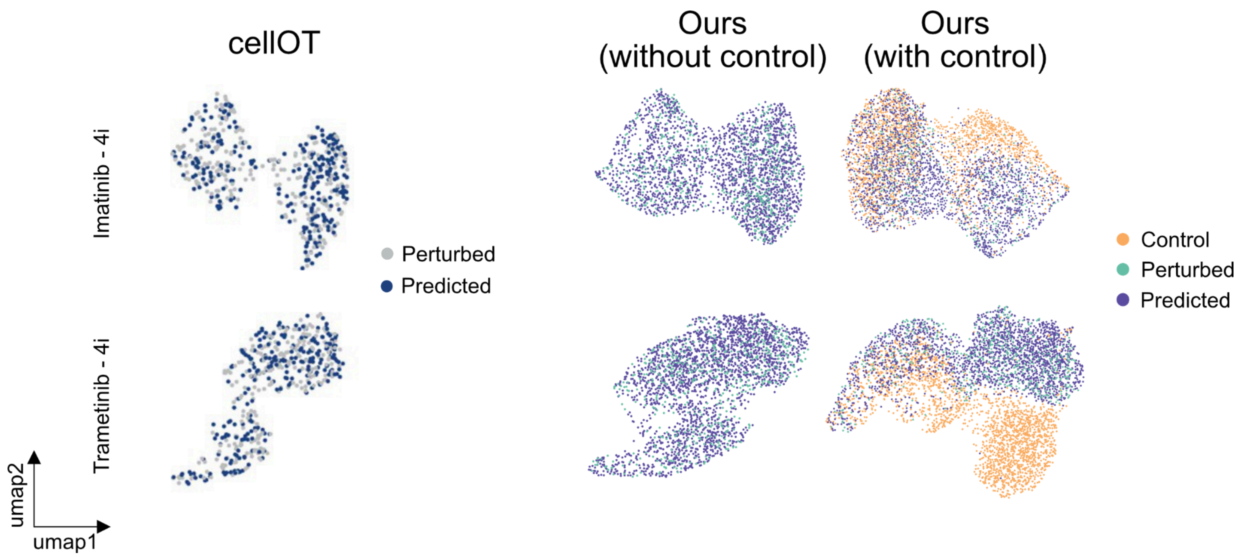


**Fig SN 5:** Comparison of results from the original cellOT publication and scArchon's reproduction on 4i data.

**scPRAM reproduction**

Fig SN6 illustrates that the predictions generated by scArchon align closely with those presented in the official scPRAM tutorial, confirming consistency between the two methods. The scPRAM tutorial used for reference is available at: https://github.com/jiang-q19/scPRAM/blob/main/Tutorial/PBMC_cross_celltype_predict.ipynb.

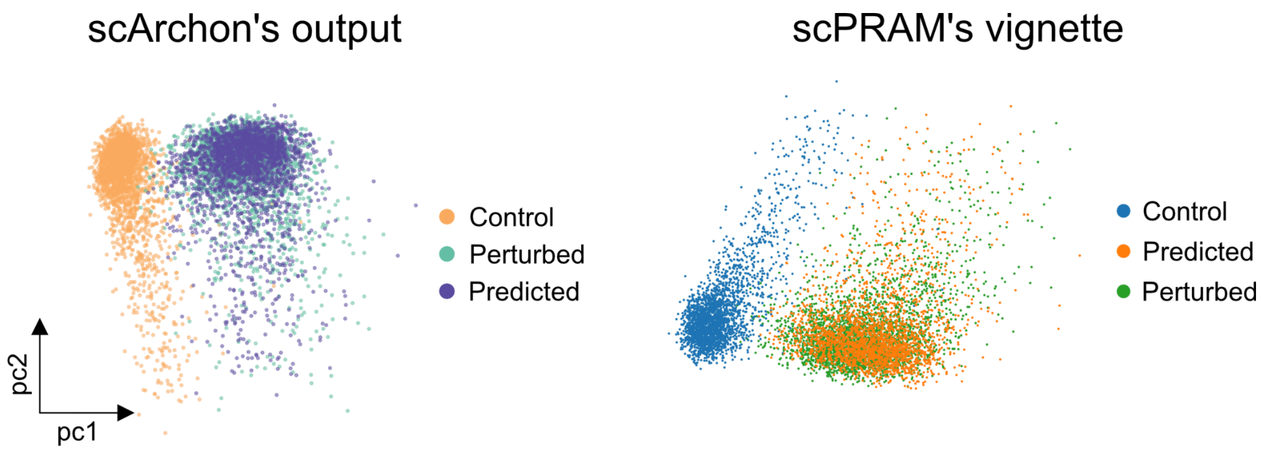


**Fig SN 6:** Comparison of prediction results from scArchon and scPRAM on CD4T cells from the Kang dataset.

### scDisInFact reproduction

The left panel shows results from scDisInFact as presented in Supplementary Fig 9 of their publication. The right panel displays the corresponding output generated using scArchon. In the scDisInFact study, the glioblastoma dataset was re-annotated to include cell type information. Our results demonstrate strong agreement with their findings. It is important to note that scDisInFact employed custom normalization and visualization techniques, which may result in visual differences. However, the underlying cell-by-gene prediction output remains consistent. The original scDisInFact implementation is available at: scDisInFact/test/test_GBM_prediction.py on their GitHub repository.

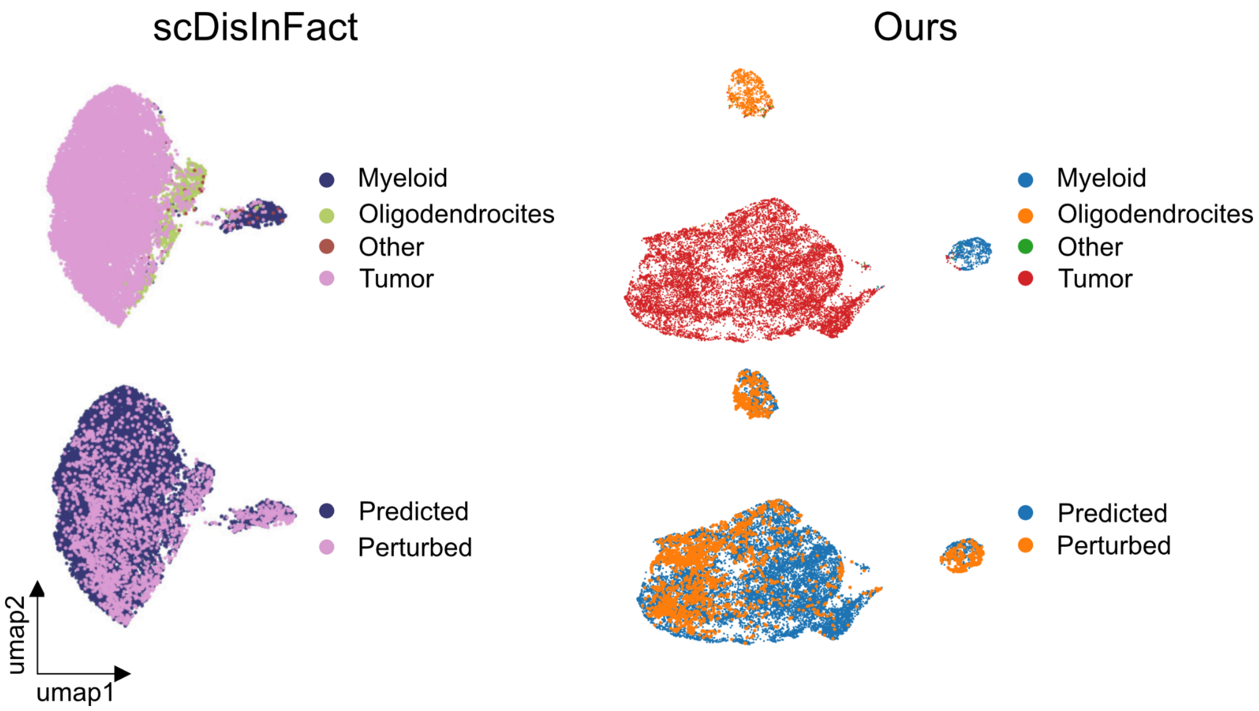


**Fig SN 7:** Comparison of scDisInFact and scArchon predictions on the glioblastoma dataset.

In their original publication, scDisInFact was compared against scGen. We aimed to verify whether our implementation of scGen within the scArchon framework would reproduce comparable results to those used in the scDisInFact study. Fig SN8 presents a side-by-side comparison of the results from scDisInFact and those generated by our scArchon implementation of scGen. It is important to note that any visual differences between these results and those shown elsewhere in scArchon stem from differences in the plotting functions used; specifically, the visualization approach in the scDisInFact paper differs from that employed in scArchon.

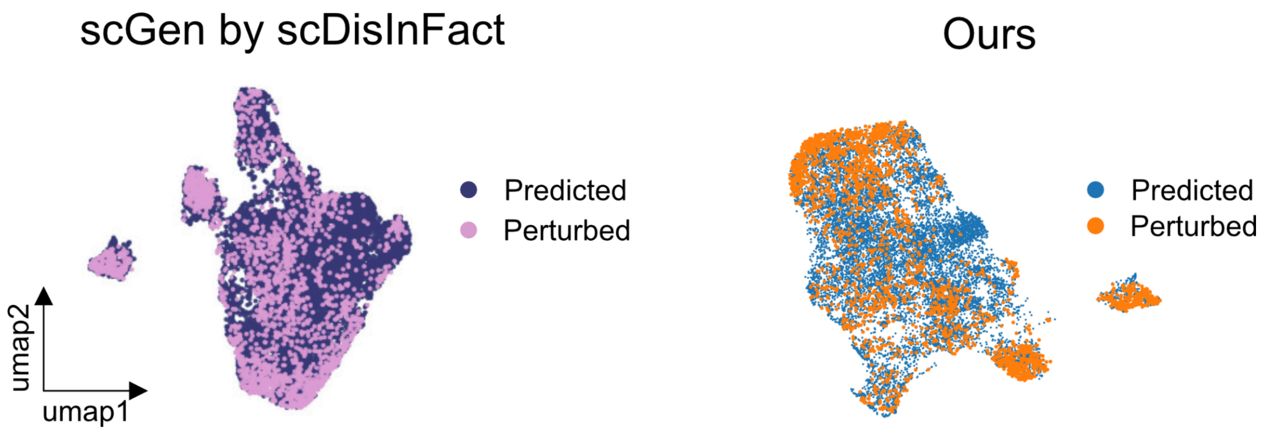


**Fig SN 8:** Comparison of scGen predictions on the glioblastoma dataset.

### CPA reproduction

As mentioned by https://github.com/theislab/cpa/issues/59, the prediction of CPA on the B cells from the Kang dataset is obtained by giving the perturbed B cells as input for the prediction, see their tutorial https://cpa-tools.readthedocs.io/en/latest/tutorials/Kang.html. Since this represents a non-sense, as we need to predict from the control and not the predicted, we changed the computation rationale by predicting on the control cells. Doing this, we observe that CPA systematically predicts the cells to map onto the control state rather than the perturbed state (Figs 2B and 2D).

### SCREEN reproduction

There is no visualisation of any results, neither in the paper nor in the supplementary. The only information we have is how to run SCREEN on CD4T. They provide Kang's adata, which is normalised (counts to 1e4 and log1p), without further visualisation of their outcome.

Li, L., You, Y., Liao, W., Fan, X., Lu, S., Cao, Y., Li, B., Ren, W., Fu, Y., Kong, J., Zheng, S., Chen, J., Liu, X., & Tian, L. (2024). A Systematic Comparison of Single-Cell Perturbation Response Prediction Models. In *bioRxiv* (p. 2024.12.23.630036). https://doi.org/10.1101/2024.12.23.630036
